# SET1/MLL complexes control transcription independently of H3K4me3

**DOI:** 10.64898/2025.12.08.692985

**Authors:** HYA Au, AT Szczurek, I de Krijger, N Kjelstrup-Osorio, AL Hughes, A Lastuvkova, M Vermeulen, RJ Klose

## Abstract

Histone H3 lysine 4 trimethylation (H3K4me3) at gene promoters is thought to play a central role in gene transcription. H3K4 methylation is deposited by the SET1 (A/B) and MLL (1-4) multi-protein complexes, but discovering how these essential enzymes shape H3K4me3 has been extremely challenging due to their multiplicity. This has also made determining whether SET1/MLL complexes control transcription through H3K4me3, or non-catalytic activities, an impenetrable problem. Here, we overcome these challenges through leveraging genome-engineering and combinatorial SET1/MLL protein depletion, integrated with genomics, proteomics, and live-cell transcription imaging. We uncover a new SET1B complex and reveal that SET1 and MLL1/2 complexes synergise to define H3K4me3 at gene regulatory elements. Unexpectedly, by decoupling SET1/MLL complex occupancy at promoters from H3K4me3, we discover they primarily control transcription independently of H3K4me3 through counteracting promoter-proximal termination and supporting transcription burst size. These discoveries reveal a new H3K4me3-independent logic for SET1/MLL-dependent control of gene transcription.

## Introduction

Gene expression patterns are instructed by transcription factors that read regulatory DNA sequences to control RNA Polymerase II (RNA Pol II) activity at gene promoters^1^. In eukaryotes, chromatin and its post-translation modifications are proposed to play central roles in shaping gene expression^2–4^. For example, chromatin-modifying complexes can localise to gene promoters and add or remove post-translational modifications to histones. While these complexes are important for gene regulation, whether they control transcription primarily through modification of histones or through their non-enzymatic properties remains a matter of vigorous ongoing debate^5–8^ and a major conceptual gap in our understanding of how DNA-encoded information is controlled to support normal cell biology and development.

This is exemplified by the SET1 and MLL family of histone H3 lysine 4 (H3K4me) methyltransferase complexes. While we know that they are essential for normal gene regulation, development, and are major targets in human disease^6,9,10^, the mechanisms they employ to regulate gene transcription have remained poorly understood. In mammals, the SET1/MLL family is composed of six large scaffolding proteins; SET1A, SET1B, MLL1, MLL2, MLL3, and MLL4, each of which contains a catalytic SET domain at its extreme C-terminus. Association of the SET domain with a complex of additional proteins called WRAD, comprising WDR5, RBBP5, ASH2L, and DPY30, enables H3K4 methylation^11–18^. *In vitro* biochemical analysis has shown that SET1A/B and MLL1/2 complexes add one (me1), two (me2), or three (me3) methyl groups to H3K4^14,19^. SET1A/B and MLL1/2 methyltransferase activity is guided to gene promoters that have CpG islands through their non-methylated DNA binding domains^20,21^. In contrast, MLL3/4 complexes primarily deposit H3K4me1/2, and can associate with distal gene regulatory elements, including enhancers, through interacting with DNA binding transcription factors^22–24^. Proteins that bind to H3K4 methylation are then thought to localise to SET1/MLL-modified chromatin to affect its structure or function, and thus regulate gene transcription^25–35^. However, emerging evidence indicates that non-catalytic activities inherent to SET1/MLL complexes may also be key to their roles in transcription regulation^36–42^. As such, despite decades of intense study, the key determinants that enable SET1/MLL complexes to regulate transcription and gene expression remain poorly understood.

Our understanding of how SET1/MLL complexes deposit H3K4 methylation and regulate gene expression in mammals originates primarily from studying gene knockouts of individual or paralogous pairs of complexes (SET1A/B, MLL1/2, or MLL3/4)^21,23,36,39,42–54^. Interestingly, these knockout approaches often yield modest influences on H3K4 methylation, presumably because other SET1/MLL complexes compensate^42,45,46^. Furthermore, analysis of transcription in SET1/MLL knockout models often yields modest or context dependent effects on gene expression, with transcriptional defects ranging from influences on initiation, pause release, and transcription elongation being reported^23,26,36,39,42,44–51,53–55^. However, because gene knockout approaches lead to the slow turnover of SET1/MLL proteins, it is impossible to determine if effects on H3K4 methylation and transcription are directly related to the disruption of SET1/MLL complex function, or a manifestation of secondary pathological consequences caused by altered cell identity or development.

To overcome the limitations of gene knockout approaches, rapid degron-based strategies have recently been employed to interrogate the function of SET1/MLL complexes. Given the challenge of simultaneously disrupting multiple SET1/MLL complexes, this has relied on depleting WRAD components (e.g. RBBP5 or DPY-30) to collectively disrupt catalysis and then interrogating how this influences H3K4 methylation and transcription^56,57^. This has led to the conclusion that H3K4me3 plays a widespread and essential role in enabling transcription, providing a histone modification-dependent logic for gene expression control by SET1/MLL complexes. However, depleting WRAD components leaves SET1/MLL proteins intact, which may still be able to control transcription through catalysis-independent mechanisms^37^. Consistent with this possibility, degron-based depletion of individual SET1/MLL proteins has indicated that there may be H3K4-methylation-independent roles for these complexes in regulating gene expression^37,41^. As such, the molecular logic through which SET1/MLL complexes control gene transcription remains poorly defined, and the extent to which this relies on H3K4me3 is still unclear.

To address these fundamental questions, we now use integrated CRISPR-based genome-engineering, degron technology, genome-wide analysis of histone modifications and nascent transcription, proteomics, and live-cell transcription imaging to dissect how SET1/MLL complexes control H3K4 methylation and transcription in mouse embryonic stem cells. We show that SET1A/B and MLL1/2 complexes synergise to deposit H3K4me3 at gene promoters and unexpectedly uncover a novel SET1B protein complex that contributes substantively to H3K4me3. Using a unique series of degron cell lines, we then discover that SET1/MLL complexes primarily control gene transcription independently of H3K4me3 and show that transcription control relies on SET1 complex occupancy. Finally, using live-cell transcription imaging we discover that SET1/MLL complexes enable gene expression by sustaining transcription burst size through counteracting stochastic promoter proximal transcription termination. Together our systematic analysis provides a new H3K4me3-independent logic for SET1/MLL complex-dependent control of gene transcription through regulating transcription burst size.

## Results

### SET1A/B and MLL1/2 complexes synergise to deposit H3K4me3

SET1A/B have been considered the central H3K4 methyltransferases that deposit H3K4me3 in mouse embryonic stem cells^53,58^. However, acute depletion of SET1A/B had surprisingly modest effects on this modification^37^, suggesting that MLL complexes may contribute substantively to H3K4me3 in this context^36,45^. To address this possibility, we set out to quantitate, for the first time under identical acute degron depletion conditions, how SET1A/B and MLL1/2 complexes control H3K4 methylation (Figures 1A and 1F). To achieve this, we used a mouse embryonic stem cell line where degradation tags (dTAGs) had been engineered into the endogenous SET1A and SET1B genes^37^ and created a distinct cell line where dTAGs were engineered into the endogenous MLL1 and MLL2 genes (Figure S1A). MLL1/2 are proteolytically cleaved after translation to create N- and C-terminal fragments^59,60^. Therefore, we engineered the dTAGs into MLL1/2 such that we could deplete their C-terminal fragments, which are required for their methyltransferase activity^59,60^. Addition of the degradation compound (dTAG13) to either SET1A/B or MLL1/2 degron cell lines resulted in rapid depletion of their respective proteins (Figures 1B and 1G).

**Figure 1:**
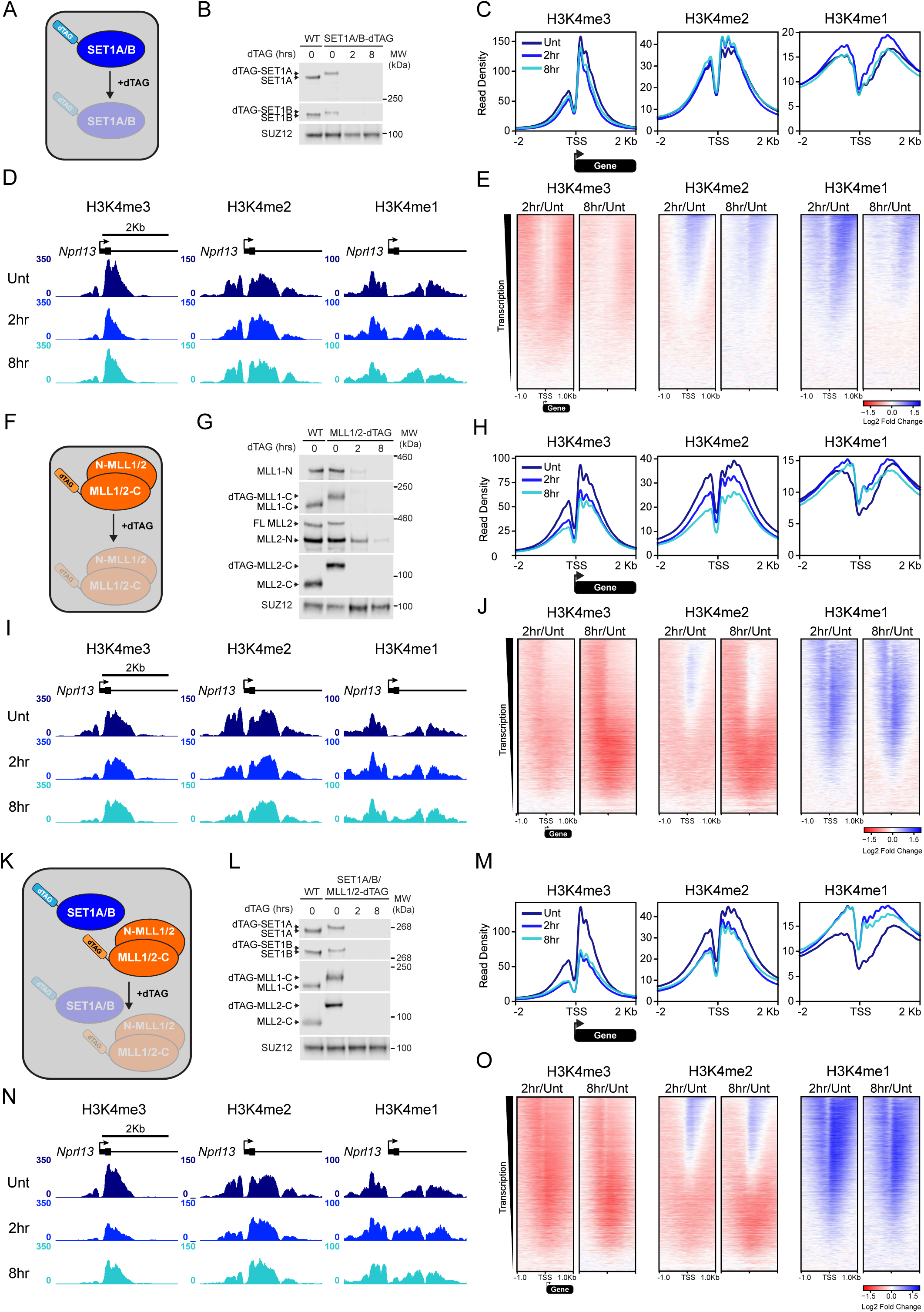
SET1A/B and MLL1/2 synergise to deposit H3K4me3. A) A schematic illustrating depletion of SET1A and SET1B. B) Western blots of SET1A and SET1B in wild type (WT) cells and SET1A/B-dTAG cells after the indicated time of dTAG13 treatment. SUZ12 is used as a loading control. C) Metaplots of average H3K4me1/2/3 ChIP-seq signal over all TSSs (n = 20,634) after SET1A/B depletion. D) Genome coverage tracks of H3K4me1/2/3 ChIP-seq signal at a representative gene (*Nprl13*) after SET1A/B depletion. E) Log2 fold change heatmaps of H3K4me1/2/3 ChIP-seq signal at all TSSs (n = 20,634) after SET1A/B depletion. Heatmaps are sorted in descending order by TT-seq signal in untreated cells from Hughes et al. 2023^37^. F) A schematic illustrating depletion of MLL1 and MLL2 C-terminal fragments (MLL1/2-C). MLL1 and MLL2 N-terminal fragments (N-MLL1/2) are also shown. G) Western blots of MLL1 and MLL2 in wild type cells and MLL1/2-dTAG cells after the indicated time of dTAG13 treatment. SUZ12 is used as a loading control. H) As per C) but after MLL1/2 depletion. I) As per D) but after MLL1/2 depletion. J) As per E) but after MLL1/2 depletion. K) A schematic illustrating co-depletion of SET1A, SET1B, MLL1, and MLL2. L) Western blots of SET1A, SET1B, MLL1, and MLL2 in wild type cells and SET1A/B:MLL1/2-dTAG cells after the indicated time of dTAG13 treatment. SUZ12 is used as a loading control. M) As per C) but after SET1A/B:MLL1/2 depletion. N) As per D) but after SET1A/B:MLL1/2 depletion. O) As per E) but after SET1A/B:MLL1/2 depletion.

We then focused on understanding how SET1A/B and MLL1/2 shape H3K4 methylation at gene transcription start sites (TSSs), given the proposed role of H3K4me3 in regulating transcription. We carried out calibrated native chromatin immunoprecipitation coupled to sequencing (cChIP-seq) for H3K4me1/2/3 at 2 and 8 hours after SET1A/B or MLL1/2 depletion (Figures 1C-1E, 1H-1J). Consistent with previous observations^37^, depletion of SET1A/B complexes had only modest effects on H3K4me3 and these occurred on nucleosomes downstream of the TSSs of highly transcribed genes (Figure 1E). In contrast, depletion of MLL1/2 caused widespread reductions in H3K4me2/3. These effects were evident on nucleosomes upstream of all TSSs, but downstream of the TSS they were limited to lowly to moderately transcribed genes (Figures 1H-1J). MLL1/2 complexes have also been implicated in shaping H3K4me3 at enhancers^21^, and we observed a clear reduction in H3K4me3 and H3K4me2 at enhancers after MLL1/2 depletion (Figure S1B). Interestingly, despite not previously having been implicated in enhancer methylation, we also observed reductions in H3K4me3 at enhancers after SET1A/B depletion, but in contrast to MLL1/2 depletion there was little effect on H3K4me2 (Figure S1C). Together this demonstrates that MLL1/2 play a central role in controlling H3K4me2/3 at TSSs, and that both MLL1/2 and SET1A/B contribute to H3K4me3 at enhancers.

Following depletion of SET1A/B and MLL1/2 individually, substantive H3K4me3 remained (Figures 1A-1J), suggesting that they may have compensatory role in defining H3K4me3 at gene regulatory elements. To explore this possibility, we developed a degron cell line where we could deplete SET1A/B and MLL1/2 simultaneously (Figures 1K and 1L). Interestingly, depletion of SET1A/B and MLL1/2 together revealed a synergistic effect on H3K4me3 on nucleosomes both upstream and downstream of TSSs across the breadth of gene transcription levels (Figure 1O) with these reductions translating into maintenance of H3K4me2 and accumulation H3K4me1. In contrast, a synergistic effect on H3K4me3 was not evident at enhancer elements (Figure S1D). This demonstrates that although SET1A/B and MLL1/2 have some inherent preference for depositing H3K4me3 at genes with different levels of transcription (Figures 1E and 1J), they synergise broadly across genes to ensure normal H3K4me3 at TSSs.

### MLL3/4 contribute only modestly to H3K4 methylation at TSSs

Simultaneous depletion of SET1A/B and MLL1/2 revealed that these complexes synergise to deposit H3K4me3. However, we were surprised that this did not result in a larger reduction in H3K4me3, given that SET1A/B and MLL1/2 are thought to be the prominent enzymes driving this modification. This suggested that additional methyltransferases may exist which contribute significantly to H3K4me3 deposition. Although MLL3/4 are primarily considered H3K4me1/2 methyltransferases that function away from TSSs, *in vitro* analysis has shown that they are also capable of installing H3K4me3 under some conditions^14^. Furthermore, we observed MLL3/4 occupancy at TSSs by cChIP-seq analysis (Figure S2A). Therefore, we reasoned that MLL3/4 may contribute to deposition of H3K4me3 at TSSs after SET1A/B:MLL1/2 depletion. To address this possibility, we developed a degron cell line in which we could deplete MLL3/4 and carried out cChIP-seq analysis of H3K4 methylation at TSSs (Figures S2B, 2A, and 2B). This revealed very little effect on H3K4me1/2 at TSSs, however there was a small but clear reduction in H3K4me3 (Figures 2C-2E). We also examined H3K4 methylation at enhancers and found that H3K4me1/2/3 were significantly reduced, in agreement with the proposed function of MLL3/4 complexes at enhancers (Figure S2C). Together this demonstrates that MLL3/4 contribute significantly to enhancer H3K4 methylation, and also modestly to H3K4me3 at TSSs.

**Figure 2:**
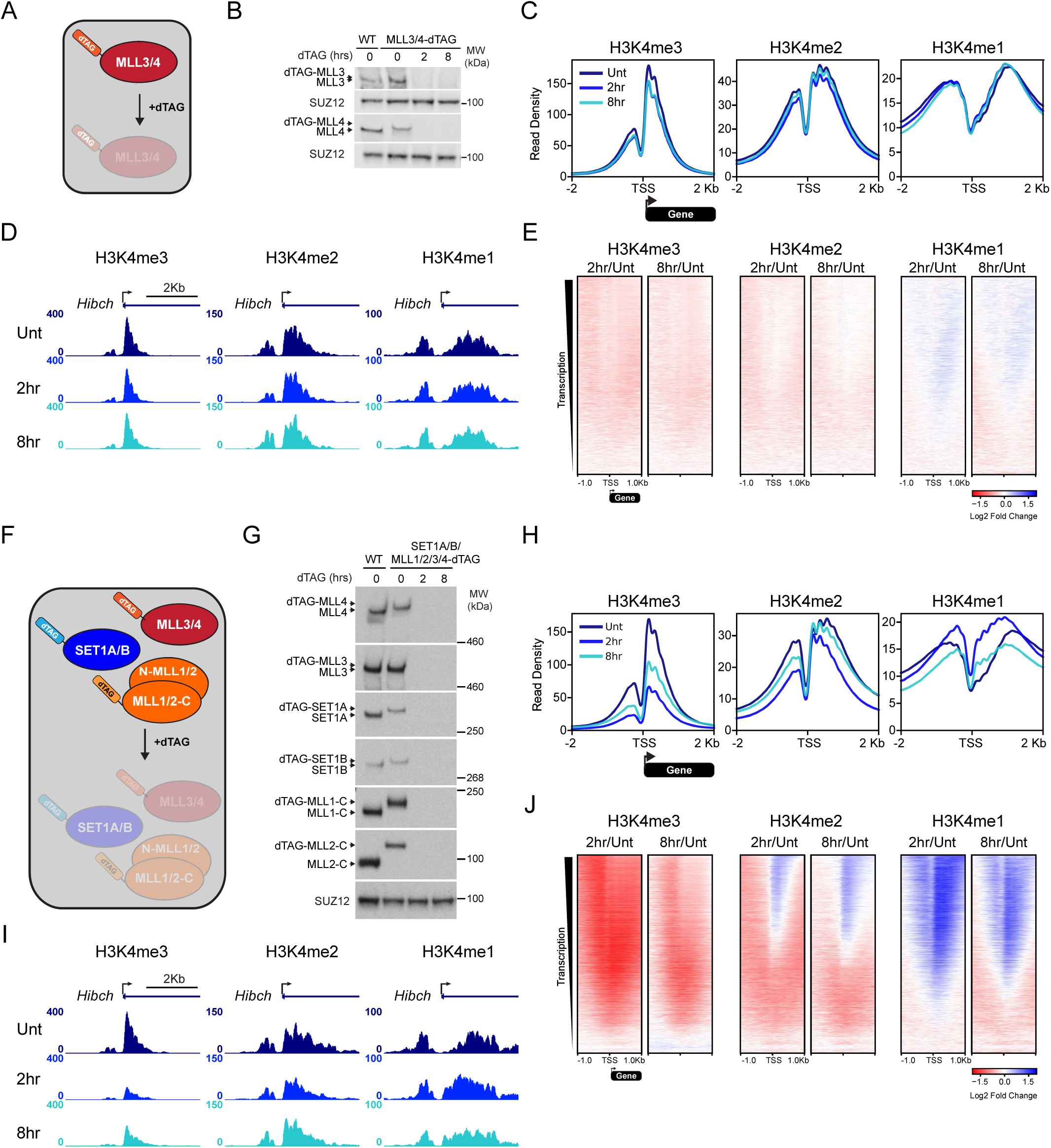
MLL3/4 contribute only modestly to H3K4 methylation at TSSs. A) A schematic illustrating depletion of MLL3 and MLL4. B) Western blots of MLL3 and MLL4 in wild type (WT) cells and MLL3/4-dTAG cells after the indicated time of dTAG13 treatment. SUZ12 is used as a loading control. C) Metaplots of average H3K4me1/2/3 ChIP-seq signal over all TSSs (n = 20,634) after MLL3/4 depletion. D) Genome coverage tracks of H3K4me1/2/3 ChIP-seq signal at a representative gene (*Hibch*) after MLL3/4 depletion. E) Log2 fold change heatmaps of H3K4me1/2/3 ChIP-seq signal at all TSSs (n = 20,634) after MLL3/4 depletion. Heatmaps are sorted in descending order by TT-seq signal untreated SET1A/B-dTAG cells as per Figure 1E. F) A schematic illustrating depletion of SET1A, SET1B, MLL1, MLL2, MLL3, and MLL4. G) Western blots of SET1A, SET1B, MLL1, MLL2, MLL3, and MLL4 in wild type cells and SET1A/B:MLL1/2/3/4-dTAG cells after the indicated time of dTAG13 treatment. SUZ12 is used as a loading control. H) As per C) but after SET1A/B:MLL1/2/3/4 depletion. I) As per D) but after SET1A/B:MLL1/2/3/4 depletion. J) As per E) but after SET1A/B:MLL1/2/3/4 depletion.

The unexpected contribution of MLL3/4 to H3K4me3 at TSSs led us to reason that MLL3/4 may synergise with SET1A/B and MLL1/2 complexes to maintain appropriate levels of H3K4me3 at TSSs. To address this possibility, we developed a degron cell line in which we could deplete all six enzymes, SET1A/B:MLL1/2/3/4, simultaneously (Figures 2F and 2G). Surprisingly, analysis of H3K4me3 at TSSs in this cell line revealed that H3K4me3 reductions were similar to those observed when SET1A/B:MLL1/2 were depleted alone (Figures 2H-2J). This indicates that although MLL3/4 localise to TSSs and can contribute to H3K4me3, they do not synergise with, nor significantly compensate for, depletion of SET1A/B:MLL1/2 in maintaining H3K4me3 at TSSs. In contrast to TSSs, the additional removal of MLL3/4 led to larger reductions in H3K4me2/3 at enhancers (Figure S2D). This indicates that MLL3/4 can in part compensate for loss of SET1A/B and MLL1/2 activity in depositing H3K4 methylation at enhancers, but not at TSSs. In fact, following the simultaneous removal of all six SET1/MLL enzymes for eight hours we observed a partial recovery of H3K4me3 at TSSs (Figures 2H-2I, S2E), suggesting, intriguingly, that an alternative H3K4 methyltransferase may compensate in depositing H3K4me3.

### Identification of a novel SET1B complex

The maintenance of H3K4me3 and its partial recovery at TSSs after prolonged depletion of SET1A/B:MLL1/2/3/4 (Figure 2H) suggested an undiscovered H3K4 methyltransferase may exist. This was unexpected as previous work using degron cell lines to deplete SET1/MLL complex-associated WRAD proteins, which are essential for catalysis, yielded a rapid and profound turnover of H3K4me3 with no apparent recovery at later time points^56,57,61^. To explore this discrepancy, we first took a biochemical approach to interrogate native SET1/MLL complexes to ensure they were efficiently depleted in the SET1/MLL degron cell line. To achieve this, we carried out size exclusion chromatography on nuclear extracts before and after SET1A/B:MLL1/2/3/4 depletion (Figures 3A and 3B). Importantly, high molecular weight complexes corresponding to SET1/MLL proteins and WRAD components were evident in untreated cells and efficiently depleted after dTAG13 treatment. Depletion of SET1A/B:MLL1/2/3/4 also resulted in an altered migration of WRAD proteins into lower molecular weight fractions, consistent with the release of core WRAD complexes and their monomeric constituents from high molecular weight SET1A/B:MLL1/2/3/4 complexes. However, interestingly, one intermediate molecular weight WRAD complex remained after SET1A/B:MLL1/2/3/4 depletion which also appeared to contain the SET1A/B complex-specific protein CFP1. This suggested that a novel complex containing WRAD proteins and CFP1 may remain after SET1A/B:MLL1/2/3/4 depletion.

**Figure 3:**
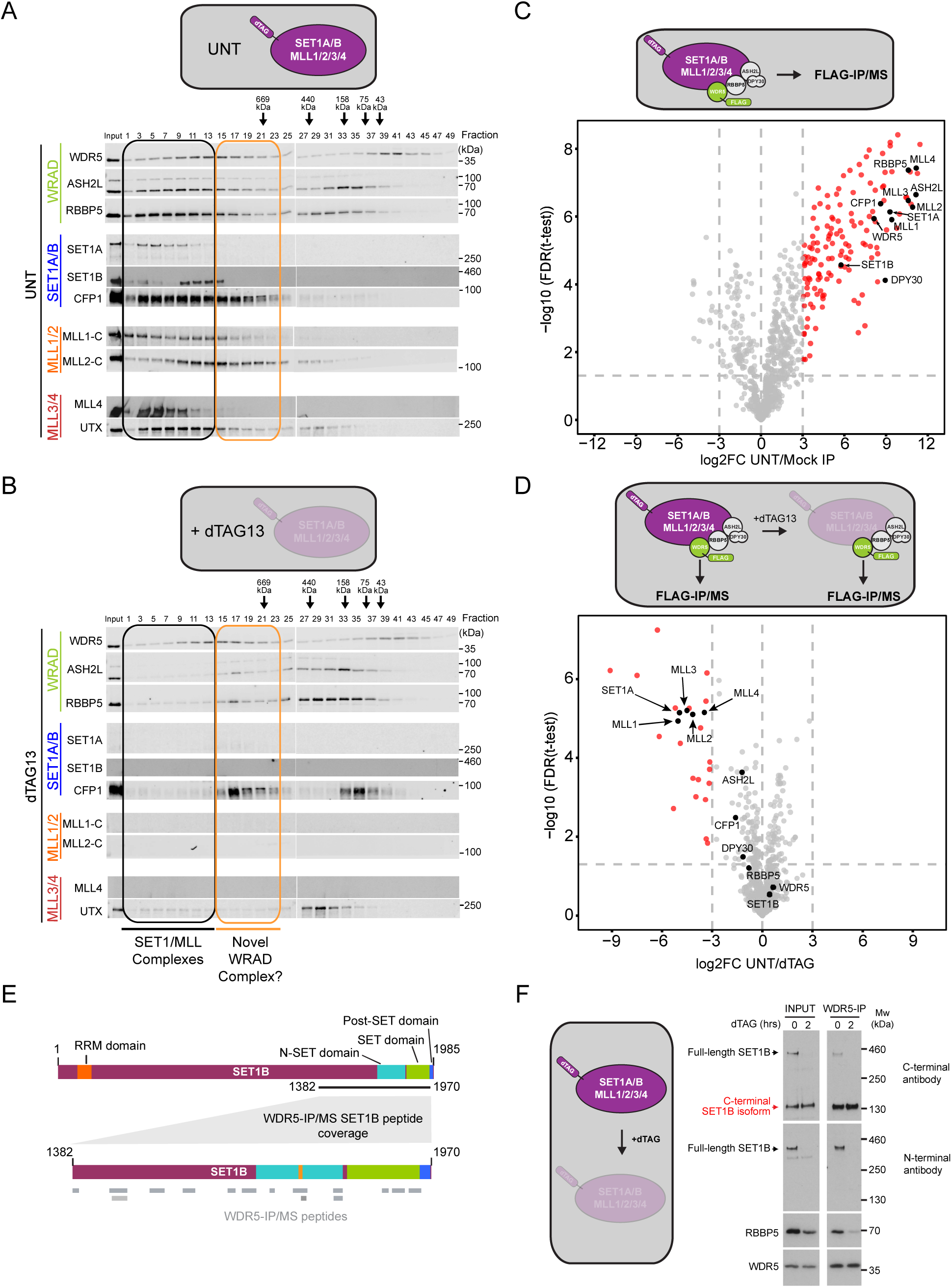
Identification of a novel SET1B complex. A) Western blots of the indicated proteins after size exclusion chromatography of nuclear extracts from untreated SET1A/B:MLL1/2/3/4-dTAG cells. B) Western blots of the indicated proteins after size exclusion chromatography of nuclear extracts from SET1A/B:MLL1/2/3/4-dTAG cells after 2 hours of dTAG13 treatment. C) Immunoprecipitation followed by mass spectrometry (IP/MS) of WDR5 in untreated SET1A/B:MLL1/2/3/4-dTAG cells. A volcano plot showing proteins identified as enriched in WDR5 purifications relative to a mock IP control. Proteins with a log2 fold change > 3 relative to the mock IP control and FDR < 0.05 are classified as statistically significant and are shown in red (n = 144). Non-significant proteins are shown in gray. SET1/MLL complex components are shown in black. D) Immunoprecipitation followed by mass spectrometry (IP/MS) of WDR5 after SET1A/B:MLL1/2/3/4-depletion. A volcano plot showing proteins identified as enriched in WDR5 purifications from 2 hour dTAG13-treated cells relative to untreated cells. Proteins with a log2 fold change > 3 relative to untreated cells and FDR < 0.05 are classified as statistically significant and are shown in red (n = 25). Non-significant proteins are shown in gray, with the exception of selected SET1/MLL complex components, which are shown in black. E) A schematic showing the location of WDR5-IP/MS peptides (grey bars) on the SET1B protein. F) Western blots of SET1B, RBBP5, and WDR5 after WDR5 immunoprecipitation from untreated and 2-hour dTAG13-treated SET1A/B:MLL1/2/3/4-dTAG cells.

Given that we had efficiently depleted SET1A/B:MLL1/2/3/4, we then sought to identify whether this intermediate molecular weight WRAD/CFP1 complex might contain a novel methyltransferase that could contribute to H3K4 methylation. To address this possibility, we used genome engineering to endogenously epitope-tag the WRAD complex component WDR5 in the SET1A/B:MLL1/2/3/4 degron cell line (Figure 3C). We then affinity-purified WDR5 and identified associated proteins by mass spectrometry (Figure 3C). This revealed known WDR5-associated protein complexes, including SET1/MLL complexes, the APC^62^, NuRD^63^, and NSL^64^, but we did not identify any methyltransferase domain-containing proteins beyond the SET1/MLL proteins themselves (Table S1). When we depleted SET1A/B:MLL1/2/3/4 and examined WDR5-associated proteins using mass spectrometry-based interaction proteomics, SET1A, MLL1/2, and MLL3/4 were significantly depleted, consistent with efficient degron-based removal of these proteins (Figure 3D). However, strikingly, we observed that SET1B, and to some extent other WRAD components and CFP1, remained enriched. Closer inspection of the enriched SET1B peptides revealed that they correspond to a C-terminal fragment of the protein (Figure 3E). This fragment encompasses the N-SET, SET, and post-SET domains that are required for WRAD association and catalysis, suggesting that a novel SET1B complex containing only the C-terminus of SET1B might exist.

Earlier western blot analysis for SET1B (Figure 1B) relied on an antibody directed against the N-terminus of SET1B which would not detect a protein comprised of only the C-terminus. To characterise this putative C-terminal SET1B protein that we identified using mass spectrometry, we used an antibody directed against the C-terminus of SET1B. By western blot analysis we observed a large molecular weight protein corresponding to full length SET1B that was efficiently depleted by dTAG13 treatment in the SET1A/B:MLL1/2/3/4 degron cell line (Figure 3F). Interestingly, with the C-terminal antibody we also observed a shorter SET1B protein of approximately 130kDa that was unaffected by dTAG13 treatment, and which could not be detected using the antibody directed against the N-terminus of SET1B. Consistent the effect on SET1B signal in these depletion experiments, the SET1B dTAG was engineered into the N-terminus of the protein and would not have been included in, nor lead to the depletion of, this shorter C-terminal form of SET1B.

To determine whether this C-terminal SET1B protein forms a WRAD complex, we immunoprecipitated WDR5 and found that the C-terminal SET1B protein associates with WDR5 independently of full length SET1B (Figure 3F). Consistent with the existence of a distinct C-terminal SET1B/WRAD/CFP1 complex, the 130kDa SET1B protein migrated with the intermediate molecular weight WRAD/CFP1 complex we identified by size exclusion chromatography (Figure S3A). To explore how this newly identified SET1B protein is produced, we examined genomics data associated with gene regulation and transcription (Figure S3B). This revealed an internal CAGE-seq peak^65^ that corresponds to a 5’ capped mRNA and accessible chromatin at a region between exons 10 and 11 of the SET1B gene, along with bi-directional nascent transcription, suggesting that an internal promoter might drive expression of a shorter SET1B protein. To examine this possibility, we carried out 5’ RACE analysis from exon 12 which yielded a defined TSS that produced an mRNA with a downstream start codon confirming the existence of a novel C-terminal SET1B isoform (Figures S3C-S3E). Therefore, our detailed systematic analysis has revealed a previously unknown SET1B complex containing WRAD proteins, CFP1, and a novel C-terminal isoform of SET1B (SET1B-C).

### The SET1B-C complex contributes centrally to H3K4me3

Based on the discovery of a new SET1B complex, we reasoned it may explain the persistent and recovering H3K4me3 we observed following depletion of full length SET1A/B:MLL1/2/3/4 (Figure 2J). Therefore, we generated a new cell line in which we engineered a dTAG into the C-terminal SET1B isoform (herein referred to as SET1B-C), allowing us to simultaneously remove all seven putative SET1/MLL methyltransferases (Figures S4A, S4B, 4A, and 4B). Importantly, the additional depletion of SET1B-C led to larger reductions in H3K4me3 at TSSs compared to depletion of only full-length SET1A/B/MLL1/2/3/4 proteins (Figures S4C-S4E). For comparison, we also created an RBBP5 degron cell line, which has previously been shown to disable the methyltransferase activity of all SET1/MLL complexes^56,57^, and carried out cChIP-seq analysis for H3K4me2/3 (Figures 4F and 4G). Importantly, following the depletion of all seven SET1/MLL complexes, the effects on H3K4me2/3 at TSSs were almost identical to those observed when RBBP5 was depleted (Figures 4C-4E, 4H-4J), demonstrating that the SET1B-C complex contributes significantly to H3K4me2/3.

**Figure 4:**
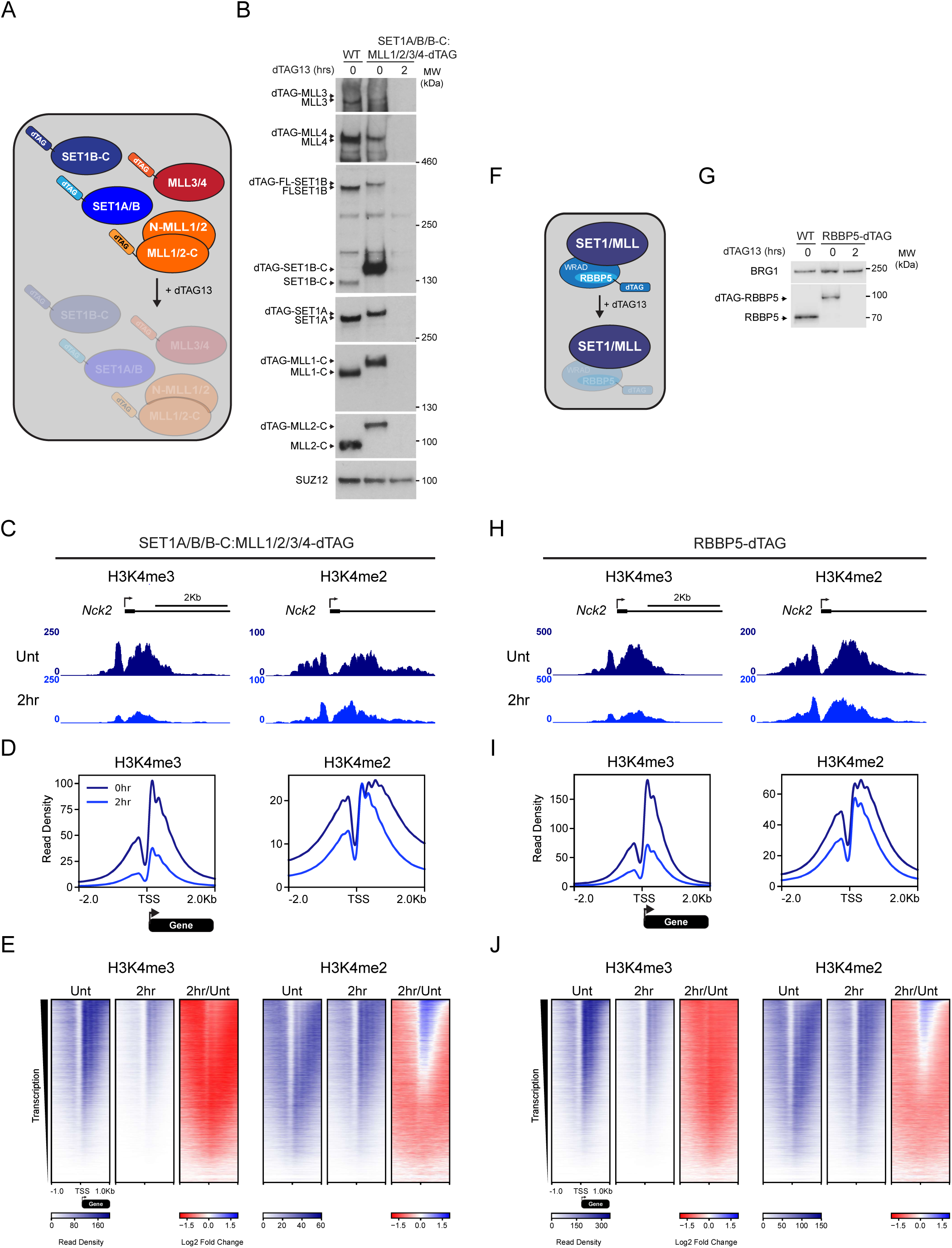
The short-SET1B complex contributes centrally to H3K4me3. A) A schematic illustrating depletion of SET1A, SET1B, SET1B-C, MLL1, MLL2, MLL3, and MLL4. B) Western blots of SET1A, SET1B, SET1B-C, MLL1, MLL2, MLL3, and MLL4 after 2 hours of dTAG13 treatment in the SET1A/B/B-C:MLL1/2/3/4-dTAG cell line. SUZ12 is used as a loading control. C) Genome coverage tracks of H3K4me2 and H3K4me3 ChIP-seq signal at a representative gene (*Nck2*) after 2 hours of dTAG13 treatment in the SET1A/B/B-C:MLL1/2/3/4-dTAG cell line. D) Metaplots of average H3K4me2 and H3K4me3 ChIP-seq signal at all TSSs (n = 20,634) after 2 hours of dTAG13 treatment in the SET1A/B/B-C:MLL1/2/3/4-dTAG cell line. E) Heatmaps of H3K4me2 and H3K4me3 ChIP-seq signal and heatmaps of log2 fold change of H3K4me2 and H3K4me3 ChIP-seq signal after 2 hours of dTAG13 treatment in the SET1A/B/ B-C:MLL1/2/3/4-dTAG cell line. All heatmaps are are centred at TSSs (n = 20,634) and sorted in descending order by TT-seq signal as in Figure 1E. F) A schematic illustrating depletion of RBBP5. G) Western blot of RBBP5 after 2 hours of dTAG13 treatment in the RBBP5-dTAG cell line. SUZ12 is used as a loading control. H) As per C) but after RBBP5-depletion. I) As per D) but after RBBP5-depletion. J) As per E) but after RBBP5-depletion.

Based on the apparent contribution of this new SET1B-C complex to H3K4me3, we reengineered our original SET1A/B and SET1A/B:MLL1/2 degron cell lines such that we could also simultaneously deplete SET1B-C (Figures S4F, S4G, S4J, and S4K). This revealed that depletion of SET1A/B/B-C caused more pronounced effects on H3K4me3 than depletion of SET1A/B alone (Figures S4H and S4I). Importantly, SET1A/B/B-C:MLL1/2 depletion caused a similar reduction in H3K4me3 to that observed when SET1A/B/B-C:MLL1/2/3/4 or RBBP5 were depleted (Figures S4L and S4M). Furthermore, both of these new cell lines showed larger reductions in H3K4me3 at enhancers than in cell lines with SET1B-C intact (Figure S4H and S4L), and removal of all seven SET1/MLL complexes caused a near complete loss of H3K4me3 at enhancers (Figure S4N). Together this demonstrates that the SET1B-C protein complex contributes significantly to H3K4 methylation, and together with full length SET1A/B and MLL1/2 complexes synergises to deposit H3K4me3 at TSS and enhancers.

### SET1/MLL complexes primarily control transcription independently of H3K4me3

Previous work using degron approaches to deplete WRAD complex components and inactivate SET/MLL complex methyltransferase activity concluded that H3K4me3 was essential for normal gene transcription^56,57^. However, studies where individual SET1/MLL complexes were disrupted also proposed there may be influences on transcription that are independent of H3K4me3^37,41^. Having now identified for the first time the full complement of SET1/MLL complexes that drive H3K4me3, and having created the tools to rapidly deplete them, we were uniquely positioned to discover how these complexes control transcription, and to determine whether this relies on H3K4me3. To do this, we took advantage of the fact that RBBP5 depletion causes major reductions in H3K4me3 but leaves the SET1/MLL proteins intact and appropriately localised to their binding sites in the genome (Figures S5A, S5B). In contrast, the simultaneous depletion of SET1A/B/B-C:MLL1/2/3/4 phenocopies the loss of H3K4me3 caused by RBBP5 depletion, but it also leads to the removal of SET1/MLL proteins from their binding sites in the genome. Therefore, a direct comparison of the effects on transcription in these two depletion contexts would allow us to determine for the first time which transcriptional effects rely on H3K4me3 and which are due to the occupancy of SET1/MLL complexes at target sites.

To address this important question, we used calibrated transient transcriptome sequencing (cTT-seq) to capture and quantify the primary effects on ongoing transcription following rapid (2 hours) depletion of SET1A/B/B-C:MLL1/2/3/4 (Figure 5A). At this time point there are major reductions in H3K4me3, but secondary or indirect transcriptional effects that could manifest from prolonged depletion are mitigated. Strikingly, following SET1/MLL complex depletion we observed ∼25% of genes (5,294) were reduced in transcription, whereas a much smaller number showed increases (Figure 5B). Furthermore, the magnitude of these effects were much larger for genes displaying decreases in transcription, indicating that SET1/MLL complexes play a widespread and important role in enabling gene transcription. We were then keen to compare these effects on transcription to those caused by RBBP5 removal where H3K4me3 is similarly reduced, but SET1/MLL complex occupancy is retained at TSSs (Figure 5E). Remarkably, RBBP5 depletion caused only minor changes in transcription, with 373 genes showing increases in transcription and 130 showing decreases (Figure 5F). This indicates that RBBP5 depletion and the turnover of H3K4me3 does not majorly influence gene transcription. Consistent with this observation, we found no correlation between changes in H3K4me3 and transcription in either depletion context (Figures 5C, 5D, 5G, 5H, and S5C). Therefore, we demonstrate that SET1/MLL complexes predominantly drive transcription through H3K4me3-independent mechanisms.

**Figure 5:**
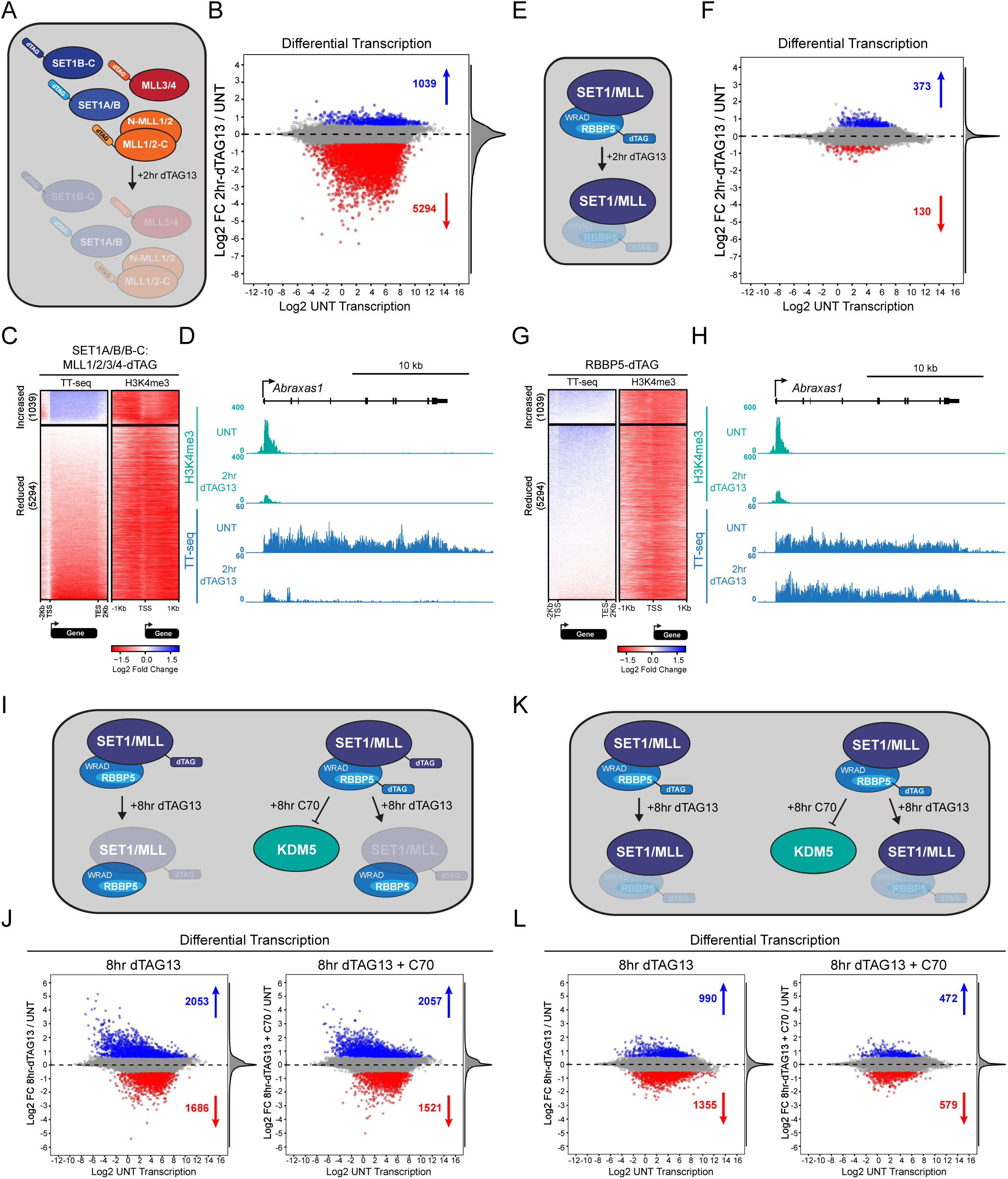
SET1/MLL complexes primarily control transcription independently of H3K4me3. A) A schematic illustrating a combined depletion of all seven H3K4 methyltransferases: SET1A, SET1B, SET1B-C, MLL1, MLL2, MLL3, and MLL4. B) An MA plot illustrating changes in TT-seq signal at all genes (n = 20,633) after depleting SET1/MLL proteins for 2 hours. The log2 fold changes (Log2 FC) are plotted on the y-axis and the log2 normalised read counts in untreated cells (Log2 UNT transcription) are plotted on the x-axis. Statistically significant increases (fold change > 1.5, *p*-adj < 0.05) are coloured in blue, and statistically significant decreases (fold change < 1.5, *p*-adj < 0.05) are coloured in red. The number of statistically significant increases and decreases are shown. C) Heatmaps illustrating the log2 fold change between 2 hour dTAG13-treated and untreated TT-seq and H3K4me3 ChIP-seq signal in the SET1A/B/B-C:MLL1/2/3/4-dTAG cell line. Heatmaps show the log2 fold change at genes that are increased (n = 1,039) or decreased (n = 5,294) in TT-seq signal after dTAG13 treatment, and sorted in descending order by the log2 fold change of the TT-seq signal. D) Genome coverage tracks showing TT-seq and H3K4me3 signal at a representative SET1/MLL-dependent gene (*Abraxas1*) after 2 hours of SET1/MLL-depletion. E) A schematic illustrating depletion of RBBP5. F) As per B) but after 2 hours of RBBP5-depletion. G) Heatmaps illustrating the log2 fold change between 2 hour dTAG13-treated and untreated TT-seq and H3K4me3 ChIP-seq signal in the RBBP5-dTAG cell line. Heatmaps show the log2 fold change at genes that are increased (n = 1,039) or decreased (n = 5,294) in TT-seq signal after dTAG13 treatment in the SET1A/B/B-C:MLL1/2/3/4-dTAG cell line. Heatmaps are sorted in descending order by the log2 fold change of the TT-seq signal in the RBBP5-dTAG cell line. H) As per D) but after 2 hours of RBBP5-depletion. I) Schematic illustrating 8 hours of SET1/MLL-depletion, with or without KDM5 inhibition using C70. J) MA plots illustrating changes in TT-seq signal at all genes (n = 20,633) after depleting SET1/MLL proteins for 8 hours, with (right) or without (left) KDM5 inhibition. The log2 fold changes (Log2 FC) are plotted on the y-axis and the log2 normalised read counts in untreated cells (Log2 UNT transcription) are plotted on the x-axis. Statistically significant increases (fold change > 1.5, *p*-adj < 0.05) are coloured in blue, and statistically significant decreases (fold change < 1.5, *p*-adj < 0.05) are coloured in red. The number of statistically significant increases and decreases are shown. K) As per I) but for RBBP5-depletion. L) As per J) but after RBBP5-depletion.

Our observations that RBBP5 depletion and turnover of H3K4me3 had little effect on transcription was unexpected given that previous work had concluded that this same perturbation profoundly affected the transcription of most active genes^56^. However, in this previous study, cTT-seq analysis was carried out at a much later time-point (8 hours) after RBBP5 depletion. To determine if this could explain the discrepancy between our findings, we carried out cTT-seq analysis in the SET1/MLL and RBBP5 degron cell lines 8 hours after depletion and included a small molecule inhibitor of the KDM5 H3K4 demethylases to examine whether transcriptional effects rely on turnover of H3K4me3 (Figure S5D). Following SET1/MLL protein depletion we observed a more complex effect on transcription, with 2,053 genes going up and 1,686 genes going down (Figures 5I and 5J). This is not surprising as the effects caused by mis-regulation of more than 5,000 genes at 2 hours will inevitably lead to secondary and compensatory effects following 8 hours of depletion. Importantly, treatment with the H3K4 demethylase inhibitor had only minor effects, demonstrating that the alterations in transcription are primarily independent of H3K4me3 (Figure 5J). After 8 hours of RBBP5 depletion, we observed some effects on transcription, but these were modest in magnitude, corresponded to roughly equal increases and decreases, and disproportionately affected lowly transcribed genes (Figures 5K, 5L, S5E). When we inhibited H3K4 demethylases, the effects on transcription of approximately half of these affected genes were significantly changed, but the reversal was incomplete (Figure S5F). Importantly, despite a near complete turnover of H3K4me3 across gene promoters following 8 hours of RBBP5 depletion, there was on average no pronounced effect on the transcription of active genes, further supporting the conclusion that H3K4me3 is not a central determinant of gene transcription (Figures S5D and S5F). Together these detailed analyses demonstrate that the depletion of H3K4me3 has minimal effects on transcription, whereas the removal of SET1/MLL complexes leads to widespread and profound reductions in transcription. Therefore, we conclude that SET1/MLL complexes primarily drive transcription independently of their effects on H3K4me3.

### SET1A/B/B-C are the main drivers of transcription

Based on the interesting observation that SET1/MLL complexes primarily regulate transcription independently of H3K4me3, we set out to discover how distinct SET1/MLL complexes shape these effects on transcription. Therefore, we first carried out cTT-seq analysis in degron cell lines where we could remove either SET1A/B/B-C or MLL1/2 complexes. Interestingly, removal of SET1A/B/B-C proteins caused profound effects on transcription, leading to a reduction in the transcription of 3,877 genes (Figure 6A). In contrast, depletion of MLL1/2 had minimal effects on transcription, despite causing much larger effects on H3K4me3 than removal of SET1A/B/B-C complexes (Figure 6B). This further reinforces that alterations in H3K4me3 are not directly linked to effects on transcription. To understand whether SET1A/B and MLL1/2 complexes may have compensatory, or even synergistic, roles in regulating transcription, we carried out cTT-seq analysis in our degron line where we can simultaneously remove SET1A/B/B-C:MLL1/2 complexes (Figure 6D). The combined depletion of SET1A/B/B-C:MLL1/2 caused only a modest increase in the number of changes beyond those observed when SET1A/B/B-C were depleted alone. The additional genes that were affected tended to be already sensitive to SET1A/B/B-C depletion, and the additional depletion of MLL1/2 led to only moderately enhanced effects on transcription (Figure S6A). Therefore, we conclude that the effects on transcription following depletion of SET1A/B/B-C:MLL1/2 are similar to those of depleting SET1A/B/B-C alone (Figure S6B).

**Figure 6:**
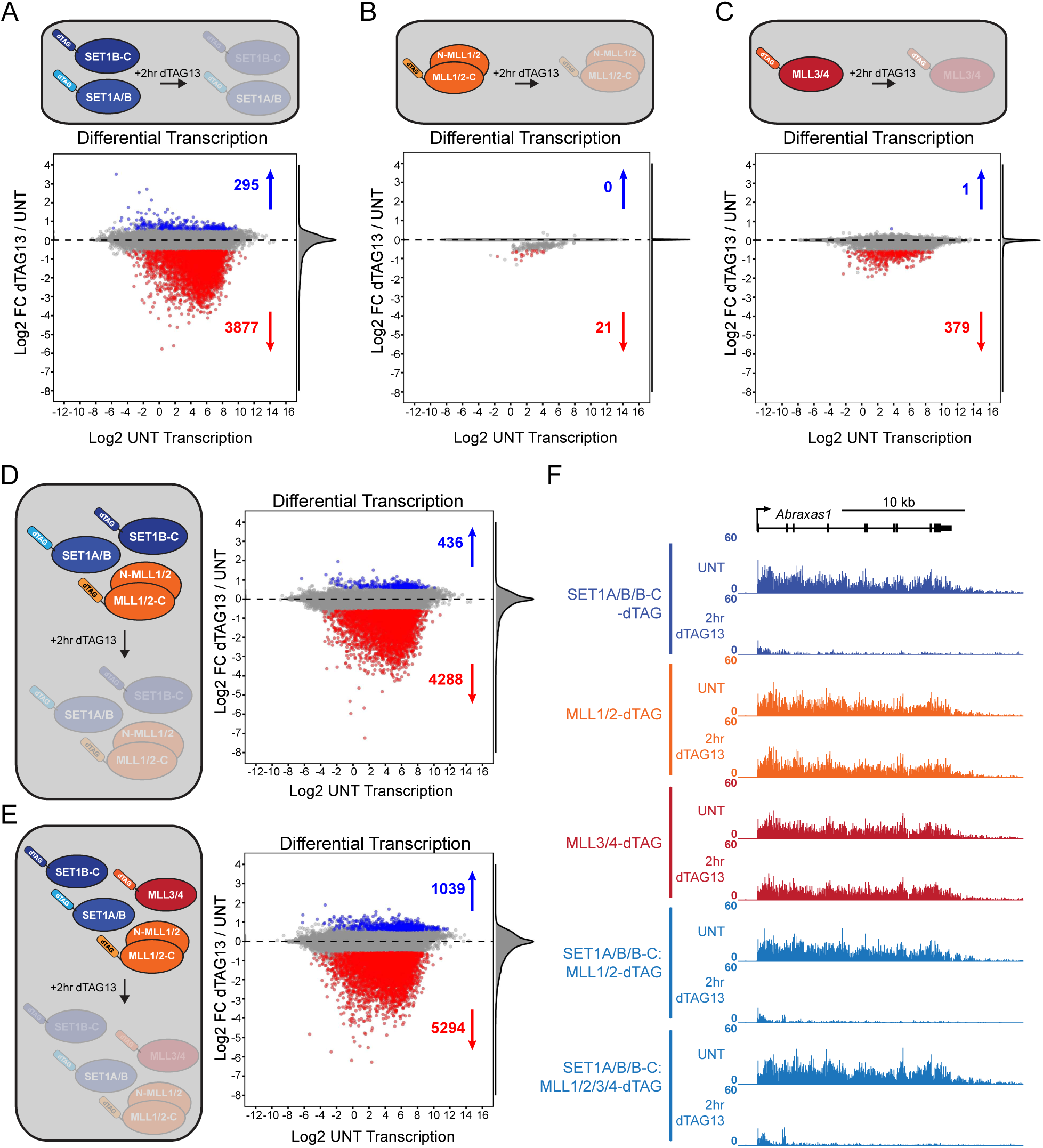
SET1A/B largely explain SET1/MLL complex dependent effects on transcription. A) An MA plot illustrating changes in TT-seq signal at all genes (n = 20,633) after depleting SET1A, SET1B, and SET1B-C proteins for 2 hours. The log2 fold changes (Log2 FC) are plotted on the y-axis and the log2 normalised read counts in untreated cells (Log2 UNT transcription) are plotted on the x-axis. Statistically significant increases (fold change > 1.5, *p*-adj < 0.05) are coloured in blue, and statistically significant decreases (fold change < 1.5, *p*-adj < 0.05) are coloured in red. The number of statistically significant increases and decreases are shown. B) As per A) but after depleting MLL1 and MLL2 for 2 hours. C) As per A) but after depleting MLL3 and MLL4 for 2 hours. D) As per A) but after depleting SET1A, SET1B, SET1B-C, MLL1, and MLL2 for 2 hours. E) As per A) but after depleting SET1A, SET1B, SET1B-C, MLL1, MLL2, MLL3, and MLL4 for 2 hours. F) Genome coverage track of TT-seq signal at a SET1/MLL-dependent gene, *Abraxas1*, after depleting the indicated proteins for 2 hours.

Given that removal of SET1A/B/B-C:MLL1/2 did not yield the extent of gene expression changes that we observed when all SET1/MLL complexes were removed, we reasoned that MLL3/4 might contribute to the increased breadth of effects on transcription. To investigate this possibility, we first carried out cTT-seq in the MLL3/4 degron cell line (Figure 6C). Interestingly, this revealed that only a few hundred genes display altered transcription and most of these effects were reductions in transcription, consistent with MLL3/4 also enabling transcription. The genes affected by MLL3/4 depletion tended to be moderately to lowly transcribed, as were the genes affected by SET1A/B/B-C depletion (Figure S6C). Therefore, we envisage that the convergence of SET1A/B/B-C:MLL3/4 activity at these genes may be important to ensure normal transcription and explain why complete SET1/MLL complex depletion affects a larger number of genes (Figure 6E). Indeed, we found that the additional genes that were uniquely reduced in transcription following complete SET1/MLL complex depletion tended to be moderately sensitive to depletion of either SET1A/B/B-C or MLL3/4 on their own (Figure S6D). We conclude that SET1A/B/B-C complexes are the primary drivers of transcription and that MLL3/4 complexes likely function with SET1A/B/B-C to enable transcription of a subset of more lowly transcribed genes.

We next examined the effect of depleting SET1/MLL complexes on enhancer transcription (Figures S6E-S6J). Like the effects on genic transcription, we found that depleting SET1A/B/B-C had the largest effects on enhancer transcription, with MLL3/4 complexes also contributing (Figures S6G-S6J). These effects appear to be independent of H3K4me3, as depletion of MLL1/2 or RBBP5 resulted in minimal changes to enhancer transcription despite causing profound reductions in H3K4me3 (Figures S6E and S6F). Therefore, we conclude that SET1 complexes are the primary effectors of transcription at both genes and enhancers, and that they primarily support transcription independently of H3K4me3 (Figure 6F).

### SET1/MLL complexes are essential to sustain transcription burst size

Our genome-wide analysis revealed that SET1/MLL complexes are essential for the transcription of thousands of genes. While cTT-seq is effective at identifying average effects on transcription at the genome scale across the cell population, it lacks the single-cell and temporal resolution that provide insight into the underlying mechanisms that ultimately shape transcription. For example, we now know from live single-cell imaging approaches that transcription is stochastic and pulsatile, occurring in bursts (ON-periods) where multiple RNA Polymerases initiate and enter productive elongation, and that these are interspersed by OFF-periods where the gene is devoid of transcription^66^. Mechanisms that regulate these features of transcription ultimately control gene expression. Therefore, to understand how SET1/MLL complexes regulate transcription we reasoned we would need to be able to examine and quantitate transcriptional features at the level of single genes in single cells.

To achieve this, we selected a SET1/MLL-dependent gene *Pank2*, and a SET1/MLL-independent gene *Hspg2*, and engineered an array of MS2 repeats into an intron of these genes to create two distinct reporter lines. These lines were also engineered to express a GFP-tagged MS2 repeat binding protein (MCP) (Figures 7A and S8A). When the MS2 labelled gene is transcribed, the resulting nascent transcript contains MS2 RNA-aptamers to which GFP-tagged MCP binds. This binding produces a discrete fluorescent focus from which we can directly quantify key parameters of the transcription cycle in live-cells in real time (Figures 7A and 7B)^67^. In addition, we further engineered these cells such that all seven SET1/MLL complexes could be depleted via the dTAG system (as above) and that RBBP5 could be independently depleted by an alternative degron system (which we call pTAG for simplicity) that is triggered by the compound PT179^68^ (Figure S8B).

**Figure 7:**
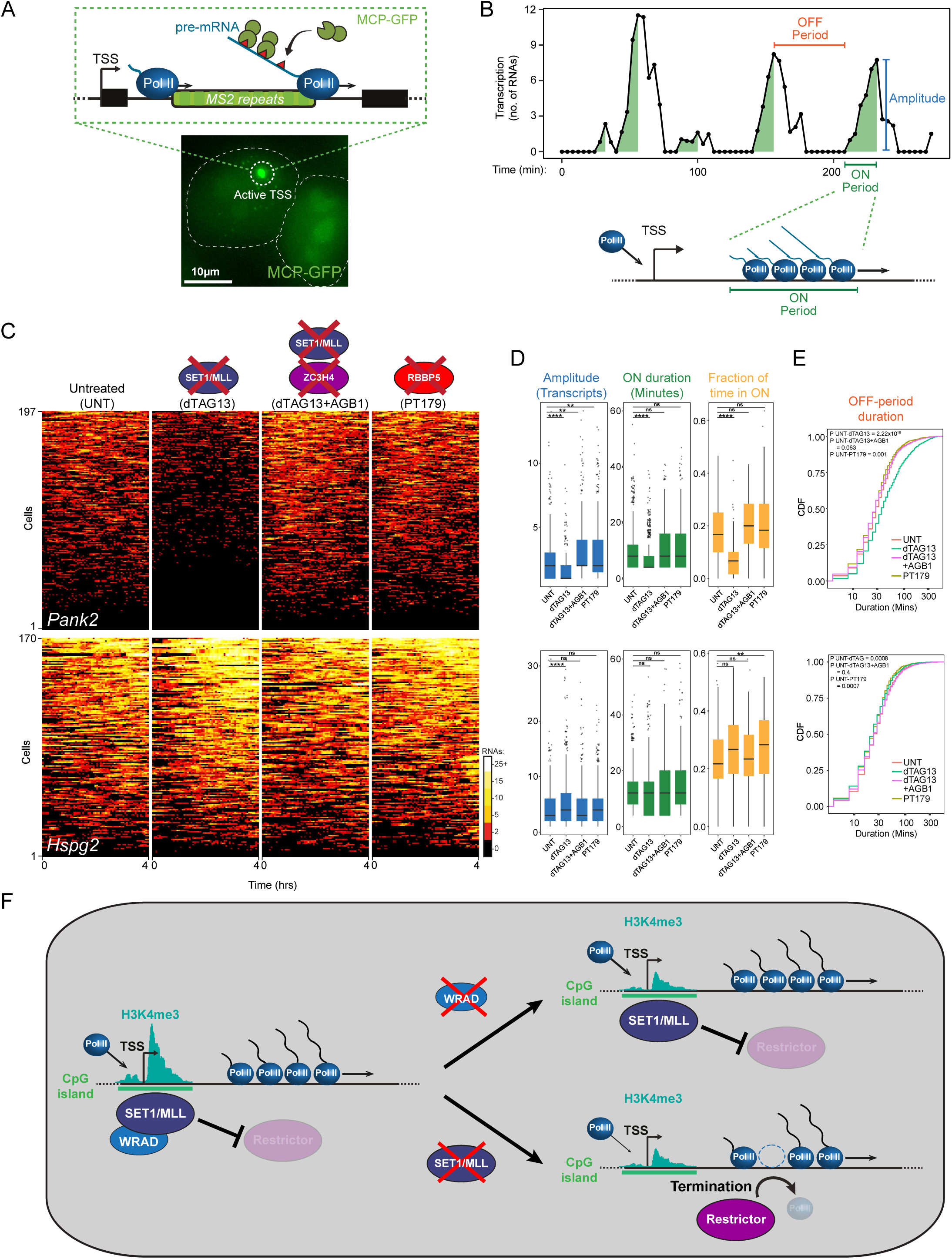
SET1/MLL complexes are essential to sustain transcription ON-period size. A) Schematic illustrating imaging of nascent transcription in living cells. Cells are engineered to constitutively express the MCP-GFP protein, and an MS2 array inserted within an intron of the gene of interest. When RNA polymerase II transcribes through the gene, the nascent transcripts produced contain the MS2 array to which MCP-GFP binds, producing a bright spot of fluorescence. Shown below is a representative image of a cell in which the active TSS produces an intense fluorescence spot. The outlines of the cells are highlighted in white. Scale bar corresponds to 10µm. B) A representative trajectory of transcriptional activity of *Pank2* over 4 hours of imaging. The key features of transcription extracted from such trajectories are shown: the ON-period duration (minutes), ON period amplitude (transcripts), and OFF-period duration (minutes). C) Kympographs of transcription trajectories from individual cells for *Pank2* and *Hspg2* after SET1/MLL-depletion (dTAG13), SET1/MLL/ZC3H4-depletion (dTAG13+AGB1), and RBBP5-depletion (PT179). The number of cells imaged over the course of 4h for each gene is indicated on the y-axis. The amplitude of transcription is colour-coded as shown on the scale bar. 3 biological replicates were imaged. D) Boxplots showing the distribution of the ON-period amplitude, ON-period duration, and the total fraction of time spent in the ON-period for each gene is shown. Each data point corresponds to an individual ON-period. Boxes represent the interquartile range (IQR) centred on the median value, with whiskers showing 1.5×IQR and outliers. Individual dots correspond to values measured for individual ON-periods, or cells for the total time spent in ON-period. *P* values were calculated using a two-sided Kolmogorov–Smirnov test, and significant *P* values are shown as asterisks: * p ≤ 0.05, ** p ≤ 0.01, *** p ≤ 0.001, and **** p ≤ 0.0001. n.s. corresponds to p > 0.05. E) Cumulative distribution function plots of the OFF-period duration. F) A cartoon illustrating the model for SET1/MLL control of transcription. When SET1/MLL complexes are physically present at CpG-island promoters, they counteract the termination activities of ZC3H4 and maintain the amplitude of the ON-period. In the absence of SET1/MLL complexes, ZC3H4 terminates transcription, resulting in a stochastic drop-out of polymerases from the Pol II convoy, thus reducing the amplitude of the ON-period. In contrast, the absence of the WRAD does not disrupt SET1/MLL occupancy, and ON-period amplitude remains intact. In both cases, H3K4me3 is depleted from the promoter.

Importantly, imaging transcription of *Pank2* or *Hspg2* revealed the expected stochastic pattern of transcription with discrete ON-periods interspersed by OFF-periods (Figure 7B). To examine how SET1/MLL complexes affect transcription, we rapidly depleted SET1/MLL complexes and first examined the transcription of the SET1/MLL-dependent gene *Pank2*. Importantly, we observed an almost immediate reduction in transcription which was apparent across the cell population (Figure 7C). In contrast, the SET1/MLL-independent gene *Hspg2* showed no such effect on transcription. We also observed similar single-cell effects on gene expression using single molecule RNA-FISH (Figure S7). We then carried out detailed quantitative analysis of key parameters of transcription for *Pank2* after SET1/MLL depletion, revealing a very clear and pronounced reduction in the ON-period amplitude and duration (Figure 7D). This led to a reduction in the fraction of time the gene spent in ON-periods and an increase in the OFF-period duration (Figure 7E). In contrast, when we depleted RBBP5 we observed no discernible effect on transcription for either *Pank2* or *Hspg2*, consistent with the effect of SET1/MLL depletion on transcription being independent of alterations to H3K4me3 (Figures 7D and 7E). This demonstrates that SET1/MLL complexes enable transcription by supporting the size of transcriptional ON-periods.

Previous models had suggested that SET1/MLL complexes may function through H3K4me3 to support transcription initiation^25,26^. In single-gene live-cell transcription imaging, effects on transcription initiation primarily manifest though altering transcription ON-period frequency^67^. However, here we find that SET1/MLL-complexes instead control the size of ON-periods and that this is independent of effects on H3K4me3. Effects on ON-period size can result from post-initiation influences on RNA Pol II; for example, transcription pause release or premature promoter-proximal termination. In analysing the effects of SET1/MLL depletion on cTT-seq signal, it is clear that most SET1/MLL-dependent genes can enter into elongation but transcription attenuates shortly thereafter, indicating that premature transcription termination may cause the observed reduction in transcription (Figure S8F). Consistent with this possibility, we previously found in genetic assays that SET1-dependent effects on transcription could be rescued by co-depletion of ZC3H4, a core component of the Restrictor transcription termination complex^37^. Therefore, we reasoned that SET1/MLL complexes may support transcription ON-period size by counteracting Restrictor-mediated transcription termination.

To test this possibility, we engineered our live-cell imaging lines so that the Restrictor component ZC3H4 could be depleted by a third degron system, the bromo-TAG (bTAG), which is triggered by the bTAG compound AGB1^69^ (Figure S8B). We first examined what effect depleting Restrictor alone had on transcription (Figures S8C-S8E). Interestingly, this revealed that transcription of SET1/MLL-dependent gene *Pank2* was slightly increased, whereas the transcription of *Hspg2* was slightly reduced. The increase in transcription of *Pank2* manifested from an increased ON-period amplitude (Figure S8D). This suggested that *Pank2* may be susceptible to Restrictor-dependent termination of transcription events within ON-periods and that SET1/MLL complexes could play a key role in counteracting this activity to support ON-period size. To directly test this possibility, we co-depleted Restrictor and SET1/MLL complexes. Remarkably, this almost completely rescued the ON-period size, as evident from the restored ON-period amplitude and duration, fraction of time the gene spends in ON-periods, and duration of OFF-periods (Figures 7C-7E). This is consistent with SET1/MLL complexes functioning to maintain ON-period size by counteracting the activity of Restrictor, which might otherwise stochastically terminate transcription within ON-periods. A model based on counteracting stochastic transcription termination is supported by the fact that we can reconstruct the observed effect on ON-period size after SET1/MLL depletion by simply calculating the frequency of lost transcriptional events and applying these as stochastic termination events to ON-periods in our unperturbed imaging data for *Pank2* (Figures S8G-S8H). In conclusion, we discover that SET1/MLL proteins support transcription burst size independently of H3K4me3 by counteracting transcription termination to enable normal gene transcription.

## Discussion

The expansion and complexity of SET1/MLL complexes in mammals had made it challenging to define how these essential systems shape H3K4 methylation and gene transcription. Here, using a unique series of degron cell lines we now discover SET1A/B and MLL1/2 complexes synergise to define H3K4me3 at gene promoters, with MLL3/4 complexes contributing only modestly (Figures 1-2). Based on our systematic dissection of H3K4me3 deposition, we discover a novel SET1B-C complex (Figure 3) and identify for the first time the complement of mammalian H3K4 methyltransferases, revealing how they shape H3K4 methylation at promoters and enhancers (Figure 4). Using this knowledge to decouple SET1/MLL complex occupancy at promoters from H3K4me3, we unexpectedly discover that SET1/MLL complexes primarily control gene and enhancer transcription independently of H3K4me3 (Figure 5), with this effect primarily relying on the activity of SET1A/B complexes (Figure 6). Using live-cell transcription imaging we then reveal that SET1/MLL complexes specifically control transcription burst size by counteracting Restrictor-dependent premature transcription termination (Figure 7).

Our discovery that SET1/MLL complexes primarily enable transcription independently of H3K4me3 was somewhat unexpected as previous work had indicated that depletion of WRAD caused pervasive effects on transcription in a manner that appeared to rely on turnover of H3K4me3^56,57^. In these studies, transcription analysis was mostly carried out after extended periods (8-12hrs) of protein depletion, which made it challenging to distinguish between the effects on transcription that were primary or secondary. Our detailed systematic analysis of SET1/MLL complexes demonstrates the importance of rapid combinatorial depletion coupled with immediate quantitative transcriptional analysis, both at the genome-scale and in live single cells, in discovering how such systems regulate transcription. We envisage our highly integrated degron-based transcriptional analysis approach will be essential for discovering how other mammalian chromatin modifying systems encoded by multiple paralogous complexes regulate histone modifications and the dynamic and cell autonomous process of gene transcription.

While we show that H3K4me3 is not required to sustain gene expression, we anticipate that it may contribute to the process of changing gene expression patterns, perhaps during events like cellular differentiation. Consistent with this, targeting of H3K4me3 to artificial reporter genes in non-permissive regions of the genome helps to activate transcription^70–74^ and catalysis by MLL2 complexes may aid in the expression of some genes during in vitro cellular differentiation^21,70^. However, under the conditions tested here, we discover that the SET1/MLL complexes function independently of H3K4me3 to enable transcription suggesting that features of these proteins beyond their methyltransferase domain must underpin this function. In support of this possibility, SET1 and MLL proteins are extremely large, comprised of several thousand amino acids, and encode multiple conserved protein domains. While some of these play important roles in targeting the complexes to their appropriate sites in the genome, other domains interact biochemically with factors that could influence the process of transcription more directly. For example, the SET1 complexes bind to a protein called WDR82 which can interact directly with RNA Polymerase II^75,76^. WDR82 is also part of the Restrictor complex, and we have previously proposed that SET1 proteins might counteract the capacity of Restrictor to influence RNA Polymerase II and ensure early elongating transcription complexes are not subject to premature transcription termination^37^. Consistent with this, in the absence of SET1 complexes, SET1-dependent genes initiate transcription and enter early elongation, but this is then rapidly terminated by Restrictor. The MLL proteins have also been proposed to function as transcription activators through directly interacting with components of the transcriptional machinery^77–79^. Therefore, it will be important in future studies to further explore how MLL proteins employ methyltransferase-independent mechanisms to control the transcription cycle and gene expression.

Transcription is much more stochastic and pulsatile than is often appreciated from the static transcriptional models that predominate in textbooks^66,80^. Yet our understanding of how stochastic transcriptional behaviors are controlled to precisely regulate gene expression remains rudimentary. Transcription factors and co-activators that acetylate histones often influence transcription burst frequency^81–83^. In contrast, we discover that depletion of SET1/MLL complexes primarily affects transcription burst size. Modulation of transcription burst size is often attributed to transcription pause release factors through their capacity to promote transcription re-initiation within bursts^84^. However, interestingly, we discover that SET1/MLL complexes instead regulate transcription burst size through a distinct mechanism that relies on counteracting the activity of the Restrictor complex that would otherwise act post-pause release^85^ to stochastically terminate individual polymerases within early elongating convoys that make up transcriptional bursts. Therefore, our systematic dissection of SET1/MLL function reveals a new H3K4me3-independent logic for gene expression control that relies on regulating transcription burst size through counteracting premature transcription termination.

## Acknowledgements

We would like to thank Jess Kelley, Tom Milne, Lars Jansen, and the Klose lab for their critical reading of the manuscript. We would also like to thank Pascal Jansen from the Vermeulen lab for technical support with mass spectrometry measurements. Work in the Klose lab is supported by the Wellcome Trust (209400/Z/17/Z). H.Y.A.A was supported by a Wellcome Trust studentship (108870/Z/15/Z).

## Data availability

The proteomics dataset generated in this study has been deposited in the ProteomeXchange Consortium via the PRIDE^86^ partner repository with the dataset identifier xxxx.

The high-thoughput data produced within this study are publicly available from the Gene Expression Omnibus (GEO) under accession number [GSEXXXXXX](https://www.ncbi.nlm.nih.gov/geo/query/acc.cgi?acc=GSE271892). Published data used in this study include TT-seq data from^37^, ATAC-seq and H3K27ac data from^87^, and CAGE-seq data from^65^.

## Materials and Methods

### Cell Culture

E14 mouse embryonic stem cells (mESCs) were cultured in Dulbecco’s minimum essential medium (DMEM, Thermofisher Scientific) supplemented with 10% fetal bovine serum (Sigma), 1x penicillin/streptomycin, 2mM L-glutamine, 1x non-essential amino acids, 0.5mM β-mercaptoethanol (Thermofisher Scientific), and 10ng/ml leukaemia inhibitory factor (LIF, produced in-house). mESCs were maintained on gelatin-coated cell culture dishes at 37°C with 5% CO2. HEK293T cells were cultured in DMEM supplemented with 10% fetal bovine serum, 1x penicillin/streptomycin, 2mM L-glutamine, 1x non-essential amino acids, 0.5mM β-mercaptoethanol and maintained on cell culture dishes at 37°C with 5% CO_2_. SG4 drosophila cells were cultured in Schneider’s Drosophila Medium (Thermofisher Scientific) supplemented with 10% heat-inactivated fetal bovine serum and 1x penicillin/streptomycin. SG4 cells were maintained adherently at 27°C.

To induce depletion of degron-tagged proteins, cells were treated with 100nM dTAG13 (Tocris Bioscience), 500nM BromoTag AGB1 (BioTechne), or 1µM of PT-179 (DC Chemicals). To inhibit KDM5 demethylases, cells were treated with 1µM KDM5-C70 (Cambridge Bioscience).

### Cell line generation

To rapidly deplete the C-terminal fragments of MLL1 and MLL2, a homozygous FKBP12^F36V^ tag was engineered into the endogenous *Kmt2a* and *Kmt2b* genes such that the tag would be inserted C-terminal to the Taspase I cleavage sites, resulting in the C-terminal fragments being N-terminally tagged after cleavage.

To rapidly deplete MLL3, a homozygous N-terminal 3xT7-2xStrepII-FKBP12^F36V^ tag was engineered into the endogenous *Kmt2c* gene. To rapidly deplete MLL4, a homozygous N-terminal 3xT7-2xStrepII - FKBP12^F36V^-eGFP tag was engineered into the endogenous *Kmt2d* gene.

To rapidly deplete RBBP5, a homozygous C-terminal 3xT7-2xStrepII-FKBP12^F36V^ tag was engineered into the endogenous *Rbbp5* gene.

To rapidly deplete the C-terminal SET1B protein, a homozygous FKBP12^F36V^ tag was inserted into exon 11 of the endogenous *Setd1b* gene such that the tag was inserted immediately after the start codon of SET1B-C, which corresponds to M1185 of the full-length protein. This also resulted in the full-length SET1B protein incorporating an internal, second copy of the FKBP12^F36V^ tag.

To rapidly deplete ZC3H4, a homozygous C-terminal 3xT7-2xStrepII-BromoTag was inserted into the endogenous *Zc3h4* gene.

To purify WDR5 from nuclear extracts, a homozygous C-terminal 3xFLAG tag was engineered into the endogenous *Wdr5* gene in the SET1A/B:MLL1/2/3/4-dTAG cell line.

CRISPR-Cas9 targeting followed by homology-directed repair was used for each genome engineering experiment. The CRISPOR online tool (http://crispor.tefor.net/crispor.py) was used to design sgRNA sequences, which were cloned into the pSptCas9(BB)-2A-Puro(PX459)-V2.0 vector, obtained from Addgene (# 91797), as previously described. The FKBP12^F36V^ tag was obtained from Addgene (# 91797) and cloned into a targeting construct vector, along with flanking homology arms of approximately 500-1000bp and any other tags as specified above. All tags included a 9 amino-acid glycine-serine linker.

Cells were transfected in one well of a 6-well plate using Lipofectamine 3000 (Thermofisher Scientific) with 0.5µg of gRNA construct and 2µg of targeting construct. Cells were plated at low density the following day and treated with 1µg/ml puromycin for 48 hours. Cells were then grown without puromycin until colonies formed, which were then picked into 96-well plates. Clones were genotyped by PCR, and homozygous clones were expanded and further characterized by immunoblotting of nuclear extracts to confirm insertion of tag.

### Calibrated native ChIP-seq

Calibrated native ChIP-seq was performed as previously described^37^. For each condition, 20 million drosophila SG4 cells were mixed with 50 million mESCs in PBS and pelleted by centrifugation at 1,500rpm for 5 minutes. Mouse-fly pellets were then resuspended in 1ml ice-cold RSB buffer (10mM Tris pH 8.0, 10mM NaCl, 3mM MgCl_2_), and nuclei were released by the addition of 28ml of ice-cold RSB buffer supplemented with 0.1% NP40. Nuclei were then pelleted by centrifugation and at 1,500g for 5 minutes at 4°C, then washed with ice-cold RSB buffer supplemented with 0.25M sucrose and 3mM CaCl_2_. Washed nuclei were pelleted as before and resuspended in 1ml of RSB buffer supplemented with 0.25M sucrose, 3mM CaCl_2_, and 1x proteinase inhibitor cocktail. Chromatin was digested with 200U of micrococcal nuclease (Thermofisher Scientific) for 5 minutes at 37°C, and 4µl of 1M EDTA was then added to stop digestion. Insoluble nuclear material was then pelleted by centrifugation at 5,000rpm at 4°C for 5 minutes, and the supernatant was then taken as the S1 fraction of chromatin. The remaining chromatin was released by resuspending the nuclear pellet with nucleosome release buffer (10mM Tris pH 7.5, 10mM NaCl, 0.2mM EDTA, 1x cOmplete protease inhibitors (Roche)) and incubating at 4°C with inversion for 1 hour. The sample was then passed through a 27G needle 5 times and insoluble material was pelleted by centrifugation at 5,000rpm for 5 minutes at 4°C. The supernatant was taken as the S2 fraction of chromatin and combined with the S1 fraction. Chromatin was then aliquoted, frozen on dry-ice, and stored at −80°C until use. DNA was purified from a small aliquot of chromatin using the QIAquick PCR purification kit (QIAGEN) and analysed on a 1% agarose gel to confirm digestion to primarily mononucleosomes.

For each ChIP reaction, 100µl of chromatin was diluted 1:10 in native ChIP incubation buffer (70mM NaCl, 10mM Tris pH 7.5, 2mM MgCl_2_, 2mM EDTA, 0.1% Triton). An additional volume of diluted chromatin was taken to use as an input sample. Diluted chromatin was centrifuged for 5 minutes at 3,000 rpm at 4°C to pellet any insoluble material. An input sample was taken from the diluted chromatin, and the remainder (1ml) was immunoprecipitated overnight at 4⁰C with anti-H3K4me1 (CST, D1A9, 5μl), anti-H3K4me2 (Invitrogen, 710796, 4μl), or anti-H3K4me3 (in-house, 1.5μl) antibody. To purify antibody-bound chromatin, 40µl blocked Protein A agarose beads (Repligen) were added to each reaction and incubated at 4°C for 1 hour with inversion. Protein A agarose beads were blocked overnight at 4°C with inversion with 1mg/ml BSA and 1mg/ml yeast tRNA in native ChIP incubation buffer. Bead-bound chromatin was washed, with gentle inversion, four times with native ChIP wash buffer (20mM Tris pH 7.5, 2mM EDTA, 125mM NaCl, 0.1% Triton) and once with TE buffer. Chromatin was eluted with 100µl elution buffer (1 % SDS, 0.1 M NaHCO_3_) at room temperature with shaking at 1,200rpm, then purified using the ChIP DNA clean and concentrator kit (Zymo Research).

For all native ChIP experiments, sequencing libraries were prepared using the NEBNext ultra II DNA Library Prep Kit for Illumina (NEB) or the NEBNext UltraExpress DNA Library Prep Kit (NEB), according to the manufacturer’s instructions.

### Calibrated Crosslinked ChIP-seq

For cChIP-seq of SET1A and MLL2, 50 million RBBP5-dTAG cells were crosslinked with 1% formaldehyde for 10 minutes at 25°C with gentle rotation. Crosslinking was then quenched with glycine added to a final concentration of 125mM, and cells were incubated at 25°C for 10 minutes with gentle rotation. Crosslinked cells were then pelleted by centrifugation for 5 minutes at 1,000g at 4°C, frozen on dry ice, and stored at −80°C until use.

For spike-in calibration, 2 million similarly crosslinked T7-RPB1 HEK293T cells were added to 50 million crosslinked mESCs. Cells were then lysed by resuspending in 1ml FA lysis buffer (50 mM HEPES pH 7.9, 150 mM NaCl, 2 mM EDTA, 0.5 mM EGTA, 0.5% NP-40, 0.1% sodium deoxycholate, 0.1% SDS, 1x protease inhibitor cocktail, 10 mM sodium fluoride) and incubation on ice for 10 minutes. Cell lysates were then sonicated on a Bioruptor Pico sonicator (Diagenode) for 30 cycles of 30 seconds on and 30 seconds off. Insoluble material was pelleted by centrifugation at 20,000g for 10 minutes at 4°C, and the supernatant was taken as chromatin. For each ChIP reaction, 100µl of chromatin was diluted 1:10 in FA lysis buffer and pre-cleared by the addition of 40µl of blocked protein A agarose beads and incubation at 4°C with inversion for 1 hour. Protein A agarose beads were blocked with 1mg/ml BSA and 1mg/ml yeast tRNA in TE buffer for 1 hour with inversion at 4°C. The appropriate antibody, anti-SET1A (abcam, ab70378, 5ul) or anti-MLL2 (CST, E3A6V, 3ul), was added to 1ml of pre-cleared chromatin and incubated overnight at 4°C with inversion. A 10% input sample was taken from the pre-cleared chromatin. Antibody-bound chromatin was obtained by incubation with 40µl of blocked protein A agarose beads for 3 hours with inversion at 4°C. Bead-bound chromatin was washed for 5 minutes with inversion at 4°C using FA lysis buffer, then high-salt FA lysis buffer (50 mM HEPES pH 7.9, 500 mM NaCl, 2 mM EDTA, 0.5 mM EGTA, 0.5% NP-40, 0.1% sodium deoxycholate, 0.1% SDS, 1x protease inhibitor cocktail, 10 mM sodium fluoride), then DOC buffer (10mM Tris-HCl pH8, 250mM LiCl, 2mM EDTA, 0.5% NP-40, 0.5% sodium deoxycholate, 10mM sodium fluoride), then twice with TE buffer. Chromatin was eluted from beads with 200µl elution buffer (1% SDS, 0.1M NaHCO_3_) at room temperature with vigorous shaking (1,200 rpm). To reverse crosslinks, NaCl was added to eluted chromatin to a final concentration of 200mM and incubated at 65C for overnight with shaking. 2µl RNase A (Sigma) was also added to remove any RNA. To remove any protein, 20µg proteinase K was added and incubated at 45°C for 1 hour with shaking. DNA was then purified using the Zymo ChIP DNA clean and concentrator kit.

For cChIP-seq of T7-tagged MLL3 and MLL4, 50 million SET1A/B/MLL1/2/3/4-dTAG cells were crosslinked with 2mM disuccinimidyl glutarate, (Thermofisher Scientific) for 45 minutes at 25°C with inversion, followed by 15 minutes of crosslinking with 1% formaldehyde (methanol-free, Thermofisher Scientific). Crosslinking was then quenched with glycine added to a final concentration of 125mM, and cells were incubated at 25°C for 10 minutes with rotation. Crosslinked cells were then pelleted by centrifugation for 5 minutes at 1,000g at 4°C, frozen on dry ice, and stored at −80°C until use.

For spike-in calibration, 2 million similarly crosslinked T7-RPB1 HEK293T cells were added to 50 million crosslinked mESCs. To isolate nuclei, cell pellets were resuspended in 10ml LB1 buffer (50mM HEPES pH7.9, 140mM NaCl, 1mM EDTA, 10% glycerol, 0.5% NP40, 0.25% Triton X100, 1x protease inhibitor cocktail) and rotated at 4°C for 10 minutes. Cells were then pelleted by centrifugation at 1000g for 5 minutes at 4°C and resuspended in 10ml of LB2 buffer (10mM Tris-HCl pH8.0, 200mM NaCl, 1mM EDTA, 0.5mM EGTA, 1x protease inhibitor cocktail). Cells were rotated at 4°C for 10 minutes, then pelleted by centrifugation at 1000g for 5 minutes at 4°C and resuspended in 1ml of LB3 buffer (10mM Tris-HCl pH8.0, 100mM NaCl, 1mM EDTA, 0.5mM EGTA, 0.5% N-laurylsarcosine, 0.1% sodium deoxycholate, 1x protease inhibitor cocktail) sonicated using a Bioruptor Pico sonicator (Diagenode) for 30 cycles of 30 seconds on and 30 seconds off. After sonication, TritonX-100 was added to a final concentration of 1%. Insoluble material was then pelleted by centrifugation for 10 minutes at 20,000g at 4°C, and the supernatant was taken as the chromatin.

For each immunoprecipitation reaction, 100µl of chromatin was diluted 1:10 with ChIP dilution buffer (1% Triton X100, 1mM EDTA, 20mM Tris-HCl, 150mM NaCl, 1x cOmplete protease inhibitors). Protein A agarose beads were blocked with 0.2 mg/ml BSA and 50 µg/ml yeast tRNA in ChIP dilution buffer for 30 minutes at 4°C with inversion. 40µl of blocked bead slurry was added to each 1ml of diluted chromatin and incubated at 4°C for 30 minutes with inversion to pre-clear. Anti-T7 antibody (CST, D9E1X, 6µl) was added to 1ml of pre-cleared chromatin and incubated overnight at 4°C with inversion. 100µl of pre-cleared chromatin was taken from each sample as an input. To capture antibody-bound chromatin, 40µl of blocked bead slurry was added to each IP reaction and incubated at 4°C with inversion for 1 hour. Beads were then washed for 4 minutes at 4°C with inversion using first 1ml of low salt buffer (1% Triton X100, 2mM EDTA, 20mM Tris-HCl, 150mM NaCl), then 1ml of high salt buffer (1% Triton X100, 2mM EDTA, 20mM Tris-HCl, 500mM NaCl), 1ml of LiCl immune complex wash buffer (250mM LiCl, 1mM EDTA, 10mM Tris-HCl, 10mg/ml sodium deoxycholate, 0.1% NP40), then twice with 1ml of TE buffer. Bead-bound chromatin was then eluted with 100µl of elution buffer (1%SDS, 0.1M NaHCO_3_), with shaking at room temperature for 30 minutes. Crosslinks were reversed by adding NaCl to a final concentration of 200mM with incubation at 65°C overnight. 2µl RNase A (Sigma) was also added. Input samples were reverse-crosslinked and RNase-treated similarly. Chromatin was then treated with 20µg of proteinase K for 1 hour at 45°C. DNA was then purified using the Zymo ChIP clean and concentrator kit.

For all crosslinked ChIP experiments, duplicate ChIP reactions were set up and purified DNA was pooled to obtain sufficient DNA for library preparation. For double-crosslinked ChIP-seq, purified ChIP and input DNA was sonicated in 50µl of elution buffer from Zymo ChIP clean and concentrator kit for 20 cycles of 30 seconds on, 30 seconds off to further fragment DNA and reduce amplification bias towards smaller fragments. For all crosslinked ChIP experiments, sequencing libraries were prepared using the NEBNext ultra II DNA Library Prep Kit for Illumina, according to the manufacturer’s instructions.

### Preparation of nuclear extracts

Cells pellets were resuspended with 10 volumes of Buffer A (10mM HEPES pH 7.9, 1.5mM MgCl_2_, 10mM KCl, 0.5mM DTT, and 0.5mM PMSF) and incubated on ice for 10 minutes. Cells were pelleted by centrifugation at 500g for 5 minutes at 4°C, then incubated with 3 volumes of Buffer A supplemented with 0.1% NP-40 at 4°C with inversion for 10 minutes to isolate nuclei. Nuclei were pelleted by centrifugation at 1,500g for 5 minutes at 4°C, then resuspended in 1 volume of Buffer C (5mM HEPES pH 7.9, 26 % glycerol, 1.5mM MgCl_2_, 0.2 mM EDTA, 1x protease inhibitor cocktail, 250mM NaCl and 0.5mM DTT). The total volume of the nuclei suspension was estimated, and 5M NaCl was added to a final concentration of 400mM. Nuclei were then incubated at 4°C with inversion for 1 hour to extract nuclear proteins. Insoluble nuclear material was then pelleted by centrifugation at 15,000g for 20 minutes at 4°C, and the supernatant was taken as the nuclear extract. Protein concentration was then measured by Bradford Assay.

### Immunoblotting

For immunoblotting of WDR5 and ASH2L, proteins were resolved by SDS-PAGE using homemade polyacrylamide gels. Proteins were then transferred onto 0.22µM nitrocellulose membranes by semi-dry transfer using the TransBlot Turbo Transfer System (BioRad), using the “high molecular weight” settings for 1 mini gel. For immunoblotting of all other proteins, proteins were resolved by SDS-PAGE using 3-8% Tris-Acetate gels. Proteins were transferred onto 0.45µM nitrocellulose membranes by wet transfer using modified Towbin’s buffer (25mM Tris, 192mM glycine, 0.01% SDS, 20% ethanol) for 22-24 hours at 25V or 90 minutes at 180V at 4°C.

Nitrocellulose membranes were then blocked with 5% milk dissolved in PBST (0.1% Tween-20 in 1X PBS) for 1 hour at room temperature with gentle agitation. Blocked membranes were incubated in the appropriate primary antibodies diluted in 5% milk-PBST overnight with gentle agitation at 4°C. Membranes were then washed three times with PBST at room temperature for 10 minutes and incubated with the appropriate secondary antibody diluted in 5% milk-PBST for 1 hour at room temperature with gentle agitation. Membranes were then washed three times with PBST at room temperature for 10 minutes and once with PBS, then imaged on a LiCor Odessey FC, or developed by enhanced chemiluminescence (ECL). For ECL, membranes were incubated with a 1:1 mixture of ECL substrate solution 1 (100mM Tris HCl, pH 8.5, 2.5mM Luminol in DMSO, 396μM p-Coumaric acid in DMSO) and solution 2 (100mM Tris HCl, pH 8.5, 5.6mM H_2_O_2_), then exposed to X-Ray film (GE Healthcare).

### Co-immunoprecipitation of WDR5 followed by western blot

Nuclear extracts were prepared from untreated and 2-hour-dTAG-treated SET1A/B:MLL1/2/3/4-dTAG cells as described above, and 1mg of nuclear extract was used for immunoprecipitation. BC150 buffer (150mM KCl, 10% Glycerol, 50mM HEPES pH 7.9, 0.5mM EDTA, 0.5mM DTT, 1x cOmplete protease inhibitors (Roche)) was added to 1mg of nuclear extract to a total volume of 550µl. Diluted nuclear extract was centrifuged at 13,000rpm for 5 minutes at 4°C to remove any insoluble precipitates, and 500µl of the supernatant was taken for immunoprecipitation. The remainder was retained as an input sample. 250U of Benzonase nuclease was added to each IP sample, followed by anti-WDR5 antibody (Bethyl, A302-429A, 5µl). Samples were incubated overnight at 4°C with inversion. Protein A agarose beads were washed twice with 1ml BC150 and blocked overnight with blocking buffer (BC150 supplemented with 1% fish skin gelatin and 0.2mg/ml BSA). To isolate antibody bound proteins, 50µl of blocked bead slurry was added to the IP samples and incubated at 4°C for 4 hours with inversion. Beads were then washed for 10 minutes with BC150 supplemented with 0.02% NP40 at 4°C with inversion six times. Bead-bound proteins were eluted by resuspending beads in 30µl of 2X SDS-PAGE loading buffer and boiled at 95°C for 5 minutes. Beads were centrifuged at 1,000g for 4 minutes at 4°C and the supernatant was analysed by immunoblotting. For immunoblotting of WDR5 and RBBP5 after WDR5-IP, VeriBlot for IP detection reagent (HRP) (Abcam) was used as the secondary antibody (1:500 dilution in 5% milk-PBST) to avoid detection of the heavy and light chains of rabbit IgG.

### Size exclusion chromatography of nuclear extracts

Nuclear extract was prepared as described above and dialysed overnight into BC200 buffer (50mM HEPES pH7.9, 200mMKCl, 10%Glycerol, 1mMDTT) at 4°C with 250U of benzonase nuclease (Merck) per mg of nuclear extract. After dialysis, samples were centrifuged at 13,000rpm for 10 minutes at 4°C to pellet any insoluble material. 1.7mg of dialysed, benzonase-treated nuclear extracts were run on a Superose 6 Increase 10/300 GL Column at flow rate of 0.2ml/min in BC200 buffer, with 250µl fractions being collected. The Superose 6 Increase 10/300 GL Column was previously calibrated with two mixes, with Ferritin (440kDa) and Conalbumin (75kDa) in mix 1, and Thyroglobulin (669kDa), Aldolase (158kDa), and Ovalbumin (43kDa) in mix 2. Alternate fractions were purified by trichloroacetic acid precipitation and the fractions were analysed by western blot.

### FLAG-immunoprecipitation of WDR5 followed by mass spectrometry

SET1A/B:MLL1/2/3/4-dTAG:WDR5-FLAG cells were treated with 100nM dTAG13 for 2 hours and harvested by scraping in ice-cold PBS supplemented with 1x cOmplete protease inhibitors (Roche). Untreated SET1A/B:MLL1/2/3/4-dTAG:WDR5-FLAG cells were also harvested as an untreated control and untreated SET1A/B:MLL1/2/3/4-dTAG cells were also harvested as a mock IP control. Cells were pelleted by centrifugation at 1,000 for 5 minutes at 4°C and stored at −80°C until use. Four biological replicates were harvested and processed as follows. Nuclear extract was prepared as described above, and 1.4mg of nuclear extract was used for FLAG purification. Nuclear extract buffer C without DTT or NaCl (5mM HEPES pH 7.9, 26 % glycerol, 1.5mM MgCl2, 0.2mM EDTA, 1x cOmplete protease inhibitors) was added to each sample to adjust the concentration of NaCl to a final concentration of 150mM. Buffer C supplemented with 150mM salt was then added to each sample to a final volume of 1.4ml. 250U of benzonase nuclease (Merck) per mg of nuclear extract was added to each sample and incubated for 30 minutes with inversion at 4°C. Insoluble material was then pelleted by centrifugation at 13,000rpm for 10 minutes at 4°C, and the supernatant was taken for immunoprecipitation. Anti-FLAG M2 affinity gel (Sigma) were washed 3 times with1ml BC150 buffer without 10% glycerol (150mM KCl, 50mM HEPES pH 7.9, 0.5mM EDTA, 0.5mM DTT, 1x cOmplete protease inhibitors), and 50µl of washed affinity gel was added to nuclear extracts and incubated at 4°C for 4 hours with inversion. Affinity gel was washed 3 times for 10 minutes at 4°C with inversion using 1ml of BC150 without 10% glycerol supplemented with 0.02% NP-40, then washed another 3 times for 10 minutes with 1ml BC150 without 10% glycerol or 0.02% NP-40. Proteins were then eluted under acidic conditions at room temperature by adding 50µl of 0.1M glycine HCl, pH 3.5, to the affinity gel, then incubating for 10 minutes with shaking at 600rpm. After centrifugation for 3 minutes at 1,000g, eluates were neutralised by transferring the supernatant into fresh tubes containing 5µl of 0.5M Tris HCl, pH 7.4, with 1.5M NaCl. The elution was repeated, and eluates were pooled and stored at −20°C until use.

For mass spectrometry analysis of WDR5-FLAG immunoprecipitants, eluates were subjected to in-solution trypsin digestion. Proteins were first denatured by adding urea to a final concentration of 4M, incubating for 10 minutes at room temperature with shaking at 650rpm. Cysteines were then reduced by adding Tris(2-carboxyethyl)phosphine to a final concentration of 10mM and incubating for 30 minutes at room temperature with no shaking. Cysteines were then alkylated by adding 2-chloroacetamide to a final concentration of 50mM and incubating for 30 minutes at room temperature in the dark with no shaking. Samples were then predigested with 1µl LysC for 2 hours at 37°C with shaking at 650rpm. The concentration of urea was then diluted to 2M using 0.1M Ammonium Bicarbonate buffer pH7.8, and CaCl_2_ was added to a final concentration of 2mM. 2µl trypsin was then added to the protein sample and incubated for 19 hours at 37°C with shaking at 800rpm. Trypsinisation was stopped by adding formic acid to a final concentration of 5%, and digested samples were centrifuged for 30mins at 12,700rpm at 20°C to remove aggregates. The supernatant was then collected and immediately desalted onto C18 columns. C18 resin was first activated by twice applying 60µl of 100% acetonitrile and centrifugation for 4 minutes at 4,000rpm at room temperature. Activated resin was then washed with 60µl of 0.1% trifluoroacetic acid, centrifuging at 4,000rpm for 4 minutes, and then 8,000rpm for 2 minutes at room temperature. The sample was then loaded onto the resin and centrifuged for 4 minutes at 4000rpm at room temperature, then centrifuged at 8000rpm for 2 minutes at room temperature. The C18 resin was then washed twice with 60µl of 0.1% trifluoroacetic acid as described above. Digested, desalted peptides were eluted twice with 60µl of 50% acetonitrile/0.1% trifluoroacetic acid, and dried overnight in a SpeedVac. Dried, digested, and desalted peptides were stored at −20°C until use.

### Liquid chromatography followed by tandem mass spectrometry

Liquid chromatography followed by tandem mass spectrometry (LC-MS/MS) was performed as previously described using an Orbitrap Exploris (Thermo)^88^. Raw mass spectrometry data were analysed with MaxQuant (version 1.6.6.0) and searched against the UniProtKB mouse proteome (Version 2017) with default settings. Identified proteins were further processed using Perseus (version 1.5.0.15). A set of four technical replicates were grouped to calculate differential proteins. Data was filtered for containing at >1 peptide identified per protein. Data were filtered for containing 4 valid values in at least 1 group, Log2 transformed, and missing values were imputated using Perseus default settings (normal distribution, width of 0.3, down shift of 1.8). Differential proteins between conditions were calculated using a two-sample t test. Data visualization was performed using R to generate volcano plots.

### Calibrated Transient Transcriptome Sequencing

Calibrated transient transcriptome sequencing (cTT-seq) was performed essentially as previously described^37,89^. 15 million mESCs were plated on a 10cm dish and treated with 100nMdTAG or 100nM dTAG + 1µM KDM5-C70 4 hours later. Cells were labelled with 500 µM 4-thiouridine (4sU, Glentham Life Sciences) for the last 15 minutes of the treatment and harvested directly into 4ml TRIzol reagent (Life Technologies). Cells harvested in TRIzol were snap frozen with ethanol and dry ice and stored at - 80°C until use. Drosophila cells were plated either in 10cm dishes (100 million cells) or 15cm dishes (300 million cells) and labelled with 500µM of 4sU for 15 minutes 3 hours later. Cells were harvested in TRIzol, snap frozen as above, and stored at −80°C until use. For spike-in calibration, 5 million drosophila cells harvested in TRIzol were added to 15 million mESCs in TRIzol. Total RNA was extracted from TRIzol using either Direct-zol RNA Miniprep Plus kit (Zymo Research) or by phenol-chloroform extraction using 15ml MaXtract High Density tubes (QIAGEN) according to the manufacturer’s instructions. RNA samples extracted with the Direct-zol Miniprep Plus kit were treated with 30U of DNase I according to the manufacturer’s instructions to remove any contaminating gDNA. Phenol-chloroform-extracted RNA samples were DNase-treated using Turbo DNA-free kit (Thermofisher Scientific). 100µg of DNase-treated total RNA was fragmented in 100µl of nuclease-free water using 20µl of 1M NaOH for 20 minutes on ice. Fragmentation was stopped by addition of 80µl of 1M Tris-HCl pH6.8 and immediately purified twice using Micro Bio-Spin P-30 gel columns (BioRad). For each experiment, an equal amount of fragmented RNA (61.54µg – 93.16µg) was biotinylated in 200µl of nuclease-free water with 50µl 0.1 mg/ml MTSEA biotin-XX linker (Biotium) and 3μl biotin buffer (833mM Tris HCl, pH 7.4, 83.3mM EDTA) for 30 minutes at room temperature in the dark. Excess biotin was removed by purifying samples twice with a 1:1 ratio of phenol/chloroform/isoamylalcohol (Thermofisher Scientific). Biotinylated RNA was enriched from an equal amount of RNA (46.53µg – 77.08µg) in 50µl of nuclease-free water using 200μl of µMACS streptavidin MicroBeads (µMACS Streptavidin Kit) and incubated with gentle agitation for 20 minutes at room temperature. Streptavidin-bound RNA was bound to µColumns and washed three times with 55 °C pull-down wash buffer (100mM Tris HCl, pH 7.4, 10mM EDTA, 1M NaCl and 0.1% Tween 20) followed by three washes with room-temperature pull-down wash buffer. Biotin-labelled RNA was eluted twice using 100µl of elution buffer (100 mM DTT in nuclease-free water) and cleaned-up using RNeasy MinElute Cleanup kit (QIAGEN), increasing the amount of 100% ethanol added to 1,050µl to capture RNA fragments smaller than 200 nucleotides.

Sequencing libraries were prepared from 20ng or 50ng of RNA using the ultra II Directional RNA library prep kit, according to the manufacturer’s instructions for rRNA depleted and FFPE RNA (NEB).

### 5’ Rapid amplification of cDNA ends (5’RACE)

5’ RACE was performed using the Template Switching RT Enzyme Mix (NEB), largely according to the manufacturer’s instructions. Total RNA was extracted from untreated SET1A/B/MLL1/2/3/4-dTAG cells using TRIzol and phenol:chloroform extraction, then DNase-treated using Turbo-DNA-free kit. 1µl of 500µg/ml Oligo(dT)_15_ Primer (Promega) and 1µl of 10mM dNTPs were added to 1µg of DNase-treated RNA, made up to a total volume of 6µl with water, and incubated at 70°C for 5 minutes to allow for annealing of the Oligo(dT)_15_ Primer to RNA. Reverse transcription and template switching was then performed by adding template switching RT buffer and RT enzyme mix to 1X, and adding the template switching oligonucleotide to a final concentration of 3.75µM in a total volume of 10µl. The reaction was incubated for 90 minutes at 42°C, then 5 minutes at 85°C. PCR amplification was then performed using a TSO-specific forward primer and 5’RACE reverse primer 1 using Q5 polymerase (NEB). The resulting PCR product was diluted 1:20 with nuclease-free water and amplified using the TSO-specific forward primer and 5’RACE reverse primer 2. The resulting amplification products were analysed on an agarose gel, gel-purified, and submitted for sanger sequencing.

### Read alignment and normalisation

For calibrated ChIP-seq, reads were aligned to the concatenated mm10+dm6 genome (for native ChIP-seq) or the mm10+hg19 genome (for crosslinked ChIP-seq) using Bowtie2^90^, with the ‘-no-mixed’ and ‘-no-discordant’ options. Only uniquely-aligned reads were retained, and PCR duplicates were removed using Sambamba^91^.

For TT-seq, reads were first aligned to mm10 rDNA sequences (GenBank: BK000964.3 and M21017.1) using Bowtie2 with ‘-very-fast’, ‘-no-mixed’ and ‘-no-discordant’ options). Reads which did not map to rDNA sequences were then mapped to the mm10 genome using STAR^92^, and any unmapped reads were re-aligned using Bowtie2 to improve mapping of intronic sequences using “–sensitive-local,” “–no-mixed” and “–no-discordant” options). Uniquely-aligned reads mapped by STAR and Bowtie2 to the mm10 genome were combined and PCR duplicates were removed using Sambamba.

For calibrated ChIP-seq, samples were internally normalized by randomly downsampling the total number of mm10 reads to reflect the number of reads aligning to the spike-in genome, essentially as previously described^93^. For native ChIP-seq, a normalisation factor was calculated using 1/(number of dm6 reads). For cross-linked ChIP-seq, variability in mixing mESC and HEK cells was considered using the ratio of hg19:mm10 reads in the input sample as previously described^93^. Briefly, the normalisation factor was adjusted by multiplying by the ratio of hg19:mm10 reads in the input sample. For each replicate, normalisation factors for each condition were divided by the largest normalisation factor for that replicate. A readcount adjustment factor was then applied such that the untreated condition in all replicates for a given histone modification or DNA-binding protein were downsampled to the same read depth, while maintaining the same ratio of spike-in normalised reads for the dTAG-treated conditions. This resulted in a downsampling factor, which was used to downsample the total number of mm10 reads using Sambamba.

For TT-seq, samples were internally normalized by randomly downsampling the total number of mm10 reads to reflect the number of reads aligning to the mouse mitochondrial genome. A readcount adjustment factor was then applied such that the untreated condition in all replicates were downsampled to the same read depth, while maintaining the same ratio of spike-in normalised reads for the dTAG-treated conditions.

Downsampled replicates were merged using Sambamba. For ChIP-seq, genome coverage files were generated using the pileup function of MACS2^94^. For TT-seq, forward and reverse strand reads were obtained using Samtools^95^, and strand-specific genome coverage files were generated using the genomeCoverageBed function from Bedtools^96^.

### Peak Calling and Annotation

For analysis of ChIP-seq and TT-seq data at genes, a previously curated list of 20,634 genes from the mm10 genome was used, in which genes were filtered to remove genes that were too short, produced similar transcripts, or had poor mappability^97^. Differential enrichment analysis of H3K4me3 ChIP-seq data was performed using DESeq2^98^ on a list of H3K4me3-associated promoters, defined as the list of promoters from the 20,634 mm10 genes described above for which the region +/-2Kb of the transcription start site overlaps with an H3K4me3 ChIP-seq peak. To obtain this list of H3K4me3-associated promoters, MACS2 was used to call peaks for H3K4me3 from all three replicates from untreated SET1A/B/MLL1/2/3/4-dTAG cells and untreated SET1A/B/MLL1/2/3/4/SET1B-C-dTAG cells. MACS2 was used without an input control, with “BAMPE” and “–-broad” specified, and a q-value cutoff of 0.05. Bedtools multiinter was used to obtain the list of H3K4me3 peaks present in all three replicates of both cell lines. This consensus peak set was filtered using Bedtools intersect with a previously-generated custom-made blacklist to remove peaks that overlapped with low-mappability and highly-repetitive regions^93^. The final list of filtered consensus H3K4me3 peaks were used to generate a list of 14,344 H3K4me3-associated promoters using Bedtools intersect.

Enhancers were previously defined using ATAC-seq and H3K27ac ChIP-seq peaks, and were filtered to remove any intragenic enhancers^87^. To additionally avoid confounding TT-seq signal from nearby genes, this list of intergenic enhancers was further filtered to remove any enhancers within 5kb of an annotated gene using Bedtools intersect to give a final list of 3,182 enhancers.

### Readcount quantification and analysis

Metaplot analysis was performed using the ComputeMatrix tool from deepTools^99^ to count the read density from genome coverage files of merged replicates across regions of interest and using plotHeatmap or plotProfile for visualisation. For metaplots of TT-seq data at genes, strand-specific read counts were obtained for sense and antisense genes, and combined using the rbind and cbind options of computeMatrixOperations to produce strand-specific metaplots. For log2 fold change heatmaps, bigwigCompare from deepTools was used to generate log2 fold change bigwig files, which were used to generate heatmaps as described above. For log2 fold change heatmaps of TT-seq data, log2 fold change bigwig files were generated from genome coverage files in which the sense and anti-sense reads were merged.

Differential transcription analysis of TT-seq was performed as described previously using DESeq2^98^. Samtools was used to obtain strand-specific read counts at the set of 20,633 genes from the mm10 genome. Samtools was also used to obtain strand-specific read counts at a set of 37 mitochondrial genes from the mm10 genome, which were used to calculate size factors for normalisation in DESeq2. The Apgelm package was used to perform shrinkage of log2 fold change values^100^. Changes were considered significant if the fold change was > 1.5 or < −1.5 and the adjusted p-value was < 0.05.

Differential enrichment analysis of H3K4me3 ChIP-seq was performed using DESeq2 analysis. multiBamSummary from deepTools was used to count reads from all three biological replicates of each condition, using the -–outRawCounts option to retain read counts. For each condition, mm10 read counts at H3K4me3-associated promoters and dm6 read counts at a custom set of 33,415 dm6 promoters were obtained. dm6 read counts were used to calculate size factors for DESeq2 analysis. Changes were considered significant if the fold change was > 2 or < −2 and the adjusted p-value was < 0.05.

Scatterplots were plotted using ggplot2, and boxplots were plotted using R. Scatterplots were coloured by density with ‘stat_density2d’ and the linear regression line was plotted using ‘geom_smooth’. Pearson’s correlation was calculated using the *cor* function. For boxplots, the box represents the interquartile range, the line inside the box represents the median, the notch represents the 95% confidence interval, and the whiskers extend to the most extreme data point within 1.5X of the interquartile range.

### Gene expression by single molecule RNA fluorescence in situ hybridization (smRNA-FISH)

To measure absolute transcript levels in single cells, the cells were prepared and imaged as described previously^101^. In brief, the cells were cultured and treated as described above. They were then trypsinised, spun down and fixed with 3.7% paraformaldehyde, and incubated in 70% ethanol/PBS. One day prior to imaging, the cells were washed with 10% formamide and 2xSSC (buffer A). Following centrifugation, the cells were incubated overnight with 200µl buffer A with 20% dextran sulfate and a set of RNA-FISH probes (25-48) labelled with Atto565 or Atto633. The following morning they were washed with buffer A, washed with 2xSSC, labelled with DAPI and Agglutinin-Alexa488, washed with PBS and resuspended on a glass slide using cell suspension : Vectashield H-1000 (Vectorlabs) at 1:1. The resulting monolayer of cells between the glass slide and the coverslip was imaged using Olympus IX83 system and cellSens software, equipped with a 63x 1.4-NA oil objective lens and a 1,200 × 1,200 px^2^ sCMOS camera (Photometrics) with a pixel size of 91.5 nm. A minimum of 300 cells were acquired per condition and per biological replicate (n=3). The data was analysed using our ThunderFISH ImageJ pipeline that allows mRNA counting in single cells: https://github.com/aleks-szczure/ThunderFISH.

### Single molecule live-cell transcription imaging and analysis

Live-cell transcription imaging with single-transcript sensitivity was performed as described previously^67^. The SET1A/B/B-C:MLL1/2/3/4/-dTAG, ZC3H4-bTAG, RBBP5-pTAG cells were engineered to express tdMCP-EGFP from a single Rosa26 locus and an array of 128 MS2 repeats was introduced into an intron of *Pank2* or *Hspg2* genes (intron 1 and 33, respectively)^102^. Prior to imaging, the medium was changed into a FluoroBrite DMEM-based medium (Thermofisher) containing dTAG-13, AGB1, or PT-179 compounds. After 40 minutes, cells on a µ-slide (IBIDI) were imaged in 3D for 4hrs at 4min time interval using Olympus IX83 system with humidified carbon dioxide chamber with a final camera pixel size of 114.4nm. Three biological replicates were acquired each containing untreated, dTAG-13, AGB1, dTAG-13+AGB1, and PT-179 conditions. Individual cells were manually cut out from the resulting 3D time-course movies and were analysed using custom-made FiJi scripts to identify nascent transcription start sites and to extract their centre of gravity, spot volume and intensity (further recalculated into single-transcript intensity as described previously^67^). The extracted spot 3D-positions were used to confirm correct spot identification in raw movies. The trajectories of transcriptional activities from individual alleles were then produced using a custom-made R script that tracks nascent transcription sites in space over time. These trajectories were then corrected for photobleaching using a GFP photobleaching standard curve. The corrected trajectories were then randomly drawn from the entire data set, grouped according to their overall transcriptional activity and plotted as kymographs with equal number of cells for all conditions including the cells that did not transcribe during the measurement time.

The resulting trajectories were then analysed using inflection points to identify local minima and maxima. This allowed identification of sections where fluorescence signal steeply raises, i.e. the times when one and more RNA PolII molecules successfully transitioned into productive elongation from a gene promoter. The identified sections of the trajectories are referred to as ON-periods (transcription bursts). The fluorescence signal increase during these periods (in single transcript intensity) is referred to as ON-period amplitude. Transcriptionally inactive sections of the trajectories correspond to OFF-periods and are defined as time intervals between the end of an ON-period and the beginning of the next ON-period. OFF-period duration is distributed as expected in a form of a long-tailed distribution. To better visualize the changes occurring primarily in its tail section, OFF-period duration was plotted in R as cumulative distribution function (CDF).

Depletion of SET1/MLLs resulted in a global decrease in transcriptional output (Figure 7, kymographs). We measured the total number of elongating RNA PolIIs in both untreated and SET1/MLL-depleted cells using our real-time live-cell transcription activity trajectories. This revealed that only 55.2% RNA PolIIs elongate across *Pank2* gene following SET1/MLL depletion. Assuming this decrease originates from a stochastic RNA PolII drop out from a gene (e.g. due to 5’-proximal transcription termination) its probability would be p_drop-out_=1 - 0.552 = 0.478. Therefore, in order to test whether RNA PolIIs are terminated stochastically, we converted the *Pank2* live-cell transcription activity trajectories from untreated condition into RNA PolII count data where individual ON-periods were represented solely by their amplitude i.e. number of RNA PolIIs (Figure S8G, mid row). We then randomly subsampled this data to simulate stochastic drop-out impacting random individual RNA PolIIs within an ON-period using p_drop-out_ (Figure S8G, bottom row). To compare the resulting “reconstructed dTAG” data to the untreated- and SET1/MLL-depleted data we have measured ON-period amplitudes and OFF-period durations.

---

**Figure S1:**
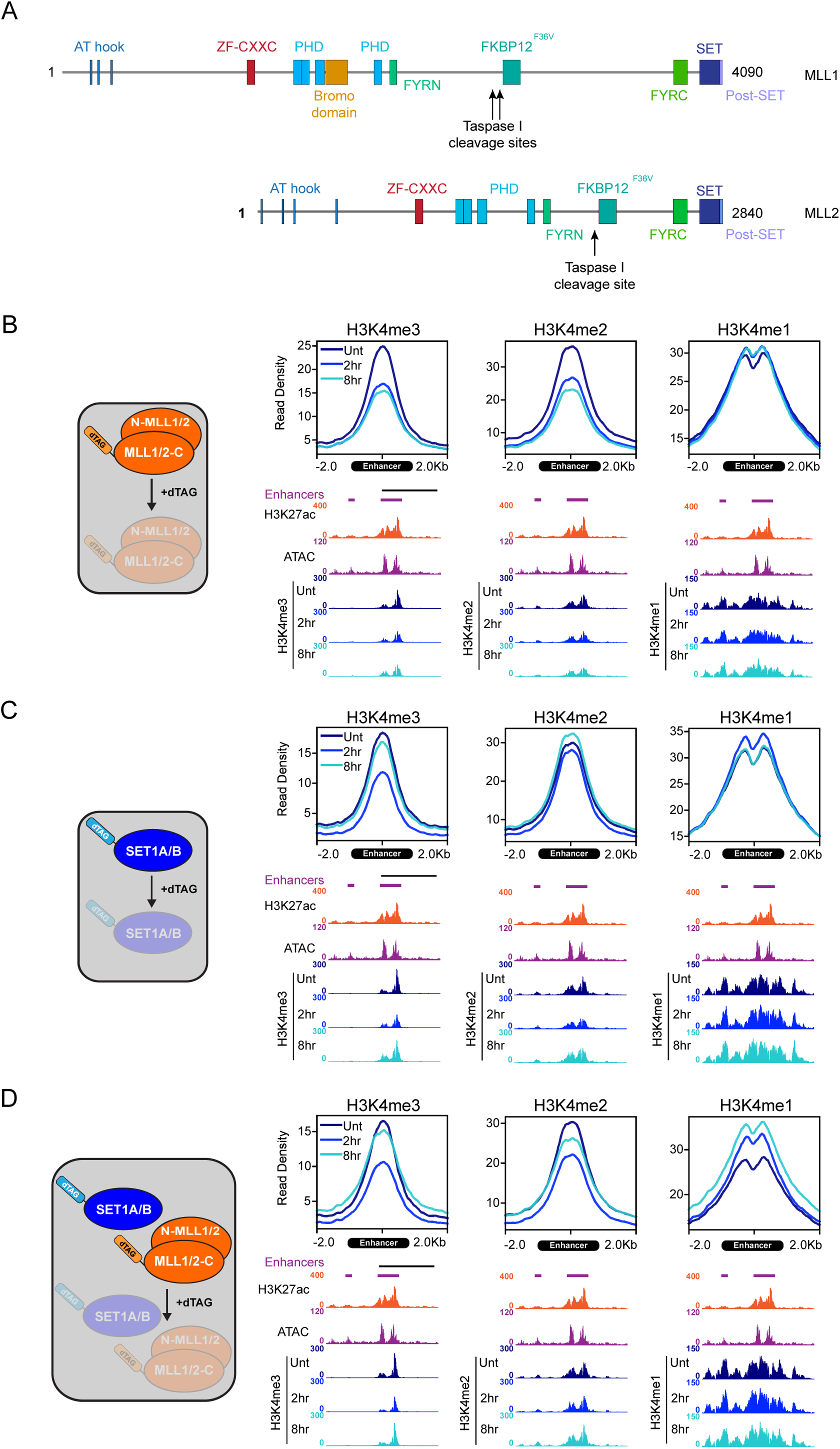
Supplementary to SET1A/B and MLL1/2 synergise to deposit H3K4me3. A) A schematic illustrating the protein domains of MLL1-dTAG and MLL2-dTAG. Taspase I cleavage sites are indicated. B) Metaplots of average H3K4me1/2/3 ChIP-seq signal in the MLL1/2-dTAG line after dTAG13 treatment at all intergenic enhancers (n=3,182) and genome coverage tracks at two enhancers at chr6: 67,105,141 – 67,106,215 and chr6: 67,110,942 – 67,114,742. Also shown are genome coverage tracks for H3K27ac ChIP-seq and ATAC-seq from E14 mESCs, both of which are from Fursova et al. 2021^87^. C) As per B) but for the SET1A/B-dTAG cell line. D) As per B) but for the SET1A/B:MLL1/2-dTAG cell line.

**Figure S2:**
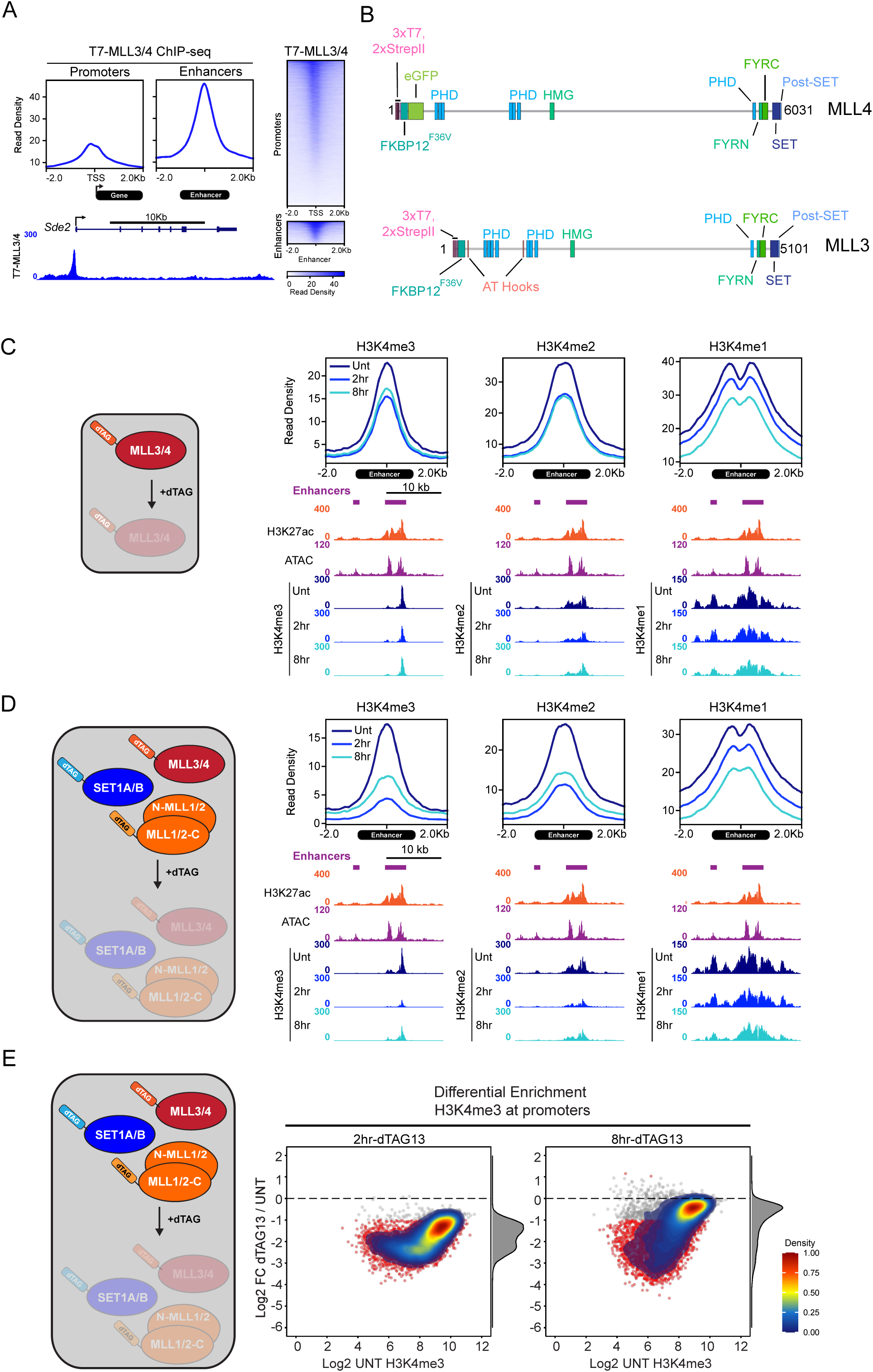
Supplementary to MLL3/4 contribute only modestly to H3K4 methylation at TSSs. A) Metaplots and heatmaps showing T7-MLL3/4 ChIP-seq signal at all TSSs (n=20,634) and intergenic enhancers (n=3,182) in untreated SET1A/B:MLL1/2/3/4-dTAG cells. Also shown is a T7-MLL3/4 ChIP-seq coverage track at a representative gene (*Sde2*). B) A schematic illustrating the protein domains of dTAG-MLL3 and dTAG-MLL4. C) Metaplots of average H3K4me1/2/3 ChIP-seq signal in the MLL3/4-dTAG line after dTAG treatment at all intergenic enhancers (n=3,182) and representative genome coverage tracks at two enhancers at chr6: 67,105,141 – 67,106,215 and chr6: 67,110,942 – 67,114,742. Also shown are genome coverage tracks for H3K27ac ChIP-seq and ATAC-seq from E14 mESCs, both of which are from Fursova et al. 2021^87^. D) As per C) but for the SET1A/B:MLL1/2/3/4-dTAG cell line. E) MA plots showing differential enrichment of H3K4me3 at all H3K4me3-associated promoters (n=14,344) between 2-hour-dTAG13-treated or 8hr-dTAG13-treated and untreated SET1A/B:MLL1/2/3/4-dTAG cells. Each dot represents an individual promoter. The log2 fold change of H3K4me3 ChIP-seq enrichment is plotted on the y-axis and the log2 normalised read count of H3K4me3 ChIP-seq in untreated cells is plotted on the x-axis. Statistically significant changes (>2 fold change, *p*-adj < 0.05) are coloured in red. A 2D density plot overlaying the data is used to illustrate the distribution of the data, with a scale bar showing the colour coding.

**Figure S3:**
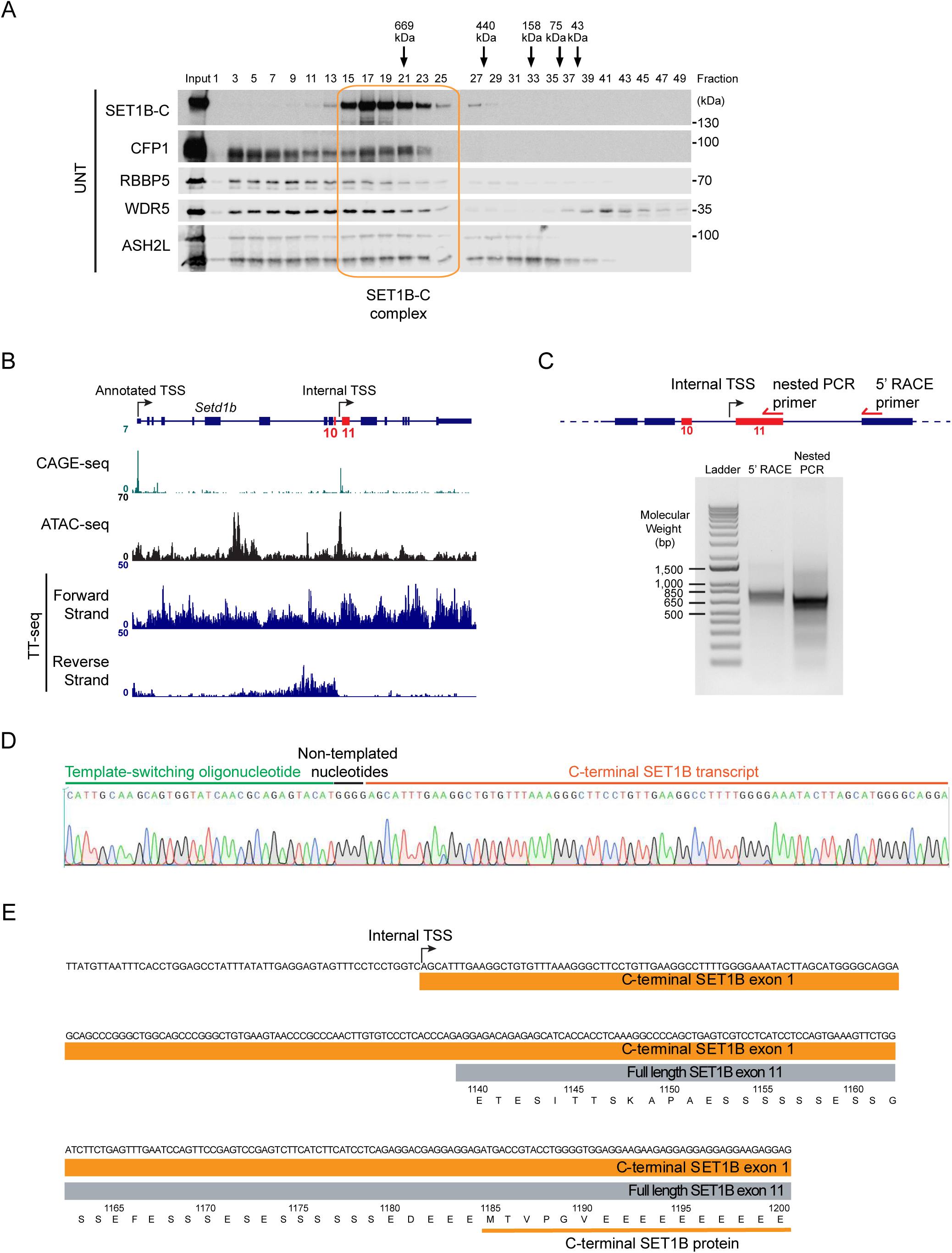
Supplementary to identification of a novel SET1B complex. A) Western blots of C-terminal SET1B (SET1B-C), CFP1, RBBP5, WDR5, and ASH2L after size exclusion chromatography of nuclear extracts from untreated SET1A/B/B-C:MLL1/2/3/4-dTAG cells. B) Genome coverage tracks of CAGE-seq, ATAC-seq, and TT-seq at the *Setd1b* gene in E14 mouse embryonic stem cells. Exons 10 and 11, flanking the internal transcription start site (TSS), are coloured and labelled in red. CAGE-seq is from Wei et al. 2020^65^, ATAC-seq is from Fursova et al. 2021^87^, and TT-seq is from Hughes et al. 2023^37^. C) 5’ rapid amplification of cDNA ends (5’ RACE) identification of the *Setd1b* internal TSS. Top – a zoomed-in view of the *Setd1b* gene. Exons 10 and 11, flanking the internal TSS, are labelled in red. The location of the primer used for 5’ RACE and subsequent nested PCR are shown (arrows not to scale). Bottom – agarose gel showing PCR amplification from the 5’RACE product using the 5’RACE reverse primer (left lane) and nested PCR on this PCR product (right lane). D) DNA sequencing traces of the nested PCR product shown in C). The sequence corresponding to the template-switching oligonucleotide used for 5’RACE is highlighted in green. Non-templated nucleotides added by the template-switching reverse transcriptase are highlighted in black. The C-terminal *Setd1b* transcript sequence is highlighted in red. E) A schematic showing the amino acid sequence of the full-length and C-terminal SET1B proteins around the internal TSS. The underlying DNA sequence and the location of the internal TSS are also shown.

**Figure S4:**
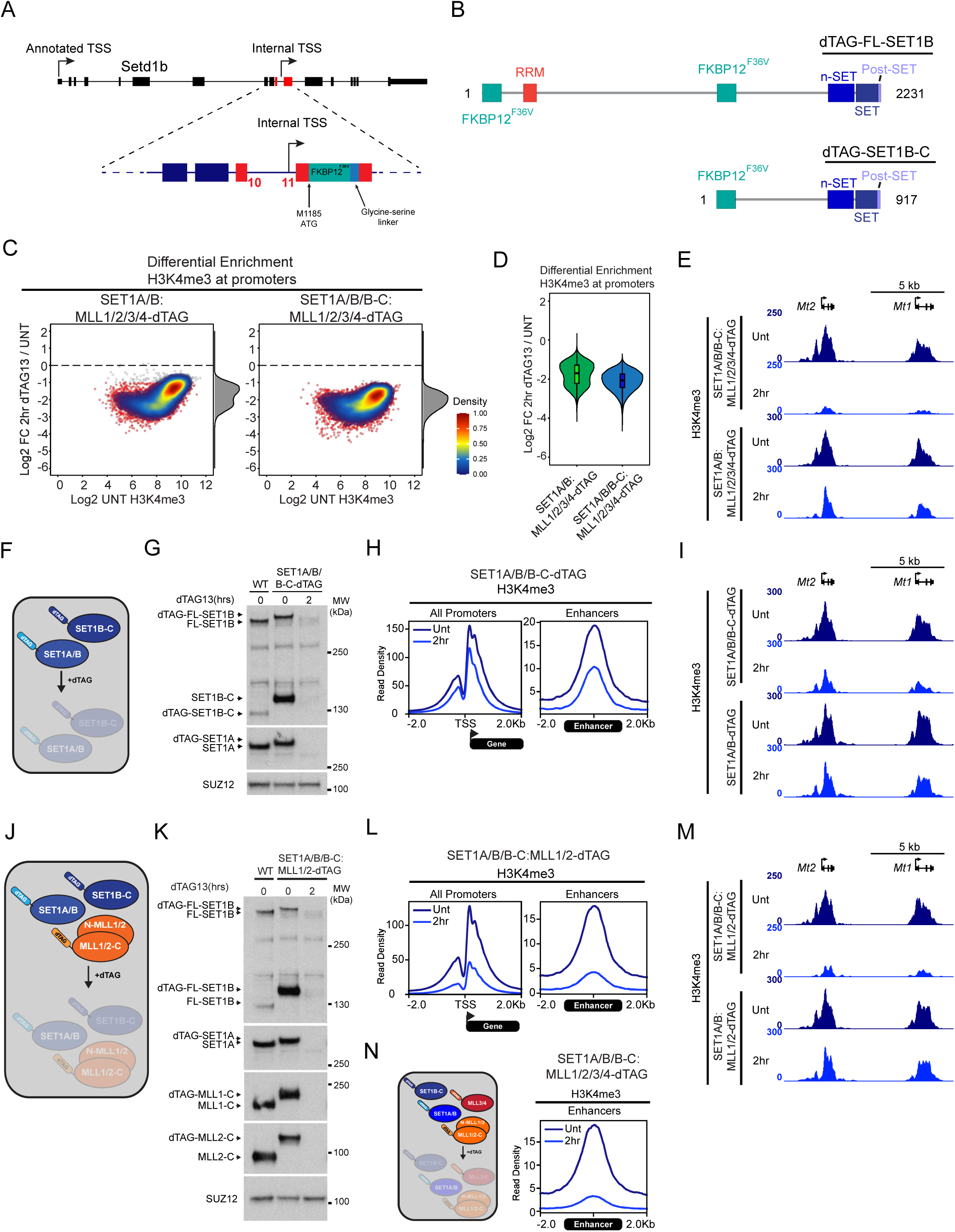
Supplementary to the short-SET1B complex contributes centrally to H3K4me3. A) A schematic illustrating genome engineering of the endogenous *Setd1b* gene to insert the FKBP12^F36V^ tag into the SET1B-C isoform. Exons 10 and 11, flanking the internal transcription start site, are highlighted in red. The start codon of the SET1B-C protein, corresponding to M1185 of the full-length SET1B protein, is indicated. B) A schematic illustrating the two dTAG-SET1B isoforms showing their protein domains. C) MA plots showing differential enrichment of H3K4me3 at all H3K4me3-associated TSSs (n=14,344) between 2hr-dTAG13 and untreated SET1A/B:MLL1/2/3/4-dTAG cells or SET1A/B/B-C:MLL1/2/3/4-dTAG cells. Statistically significant changes (>2 fold change, *p*-adj < 0.05) are coloured in red. A 2D density plot overlaying the data is used to illustrate the distribution of the data, with a scale bar showing the colour coding. D) Violin plots showing differential enrichment of H3K4me3 at all H3K4me3-associated promoters (n=14,344) between 2hr-dTAG13 and untreated SET1A/B:MLL1/2/3/4-dTAG cells or SET1A/B/B-C:MLL1/2/3/4-dTAG cells. E) Genome coverage tracks of H3K4me3 at two representative genes (*Mt2, Mt1*) after 2 hour dTAG13 treatments in the SET1A/B/B-C:MLL1/2/3/4-dTAG and SET1A/B:MLL1/2/3/4-dTAG cells. F) A schematic illustrating depletion of SET1A, SET1B, and SET1B-C. G) Western blots of SET1A, SET1B, and SET1B-C after 2 hours of dTAG13 treatment in the SET1A/B/B-C-dTAG cell line. SUZ12 is used as a loading control. H) Metaplot of average H3K4me3 ChIP-seq signal over all TSSs (n=20,634) and intergenic enhancers (n=3,182) after 2 hours of dTAG13 treatment in the SET1A/B/B-C-dTAG cell line. I) As per (E) but for the SET1A/B/B-C-dTAG and SET1A/B -dTAG cell lines. J) A schematic illustrating depletion of SET1A, SET1B, SET1B-C, MLL1, and MLL2. K) Western blots of SET1A, SET1B, SET1B-C, MLL1, MLL2 after 2 hours of dTAG13 treatment. SUZ12 is used as a loading control. L) As per (H) but for the SET1A/B/B-C:MLL1/2-dTAG cell line. M) As per (E) but for the SET1A/B/B-C:MLL1/2-dTAG and SET1A/B:MLL1/2-dTAG cell lines. N) Metaplot of average H3K4me3 ChIP-seq signal over all intergenic enhancers (n=3,182) after 2 hours of dTAG13 treatment in the SET1A/B/B-C:MLL1/2/3/4-dTAG cells.

**Figure S5:**
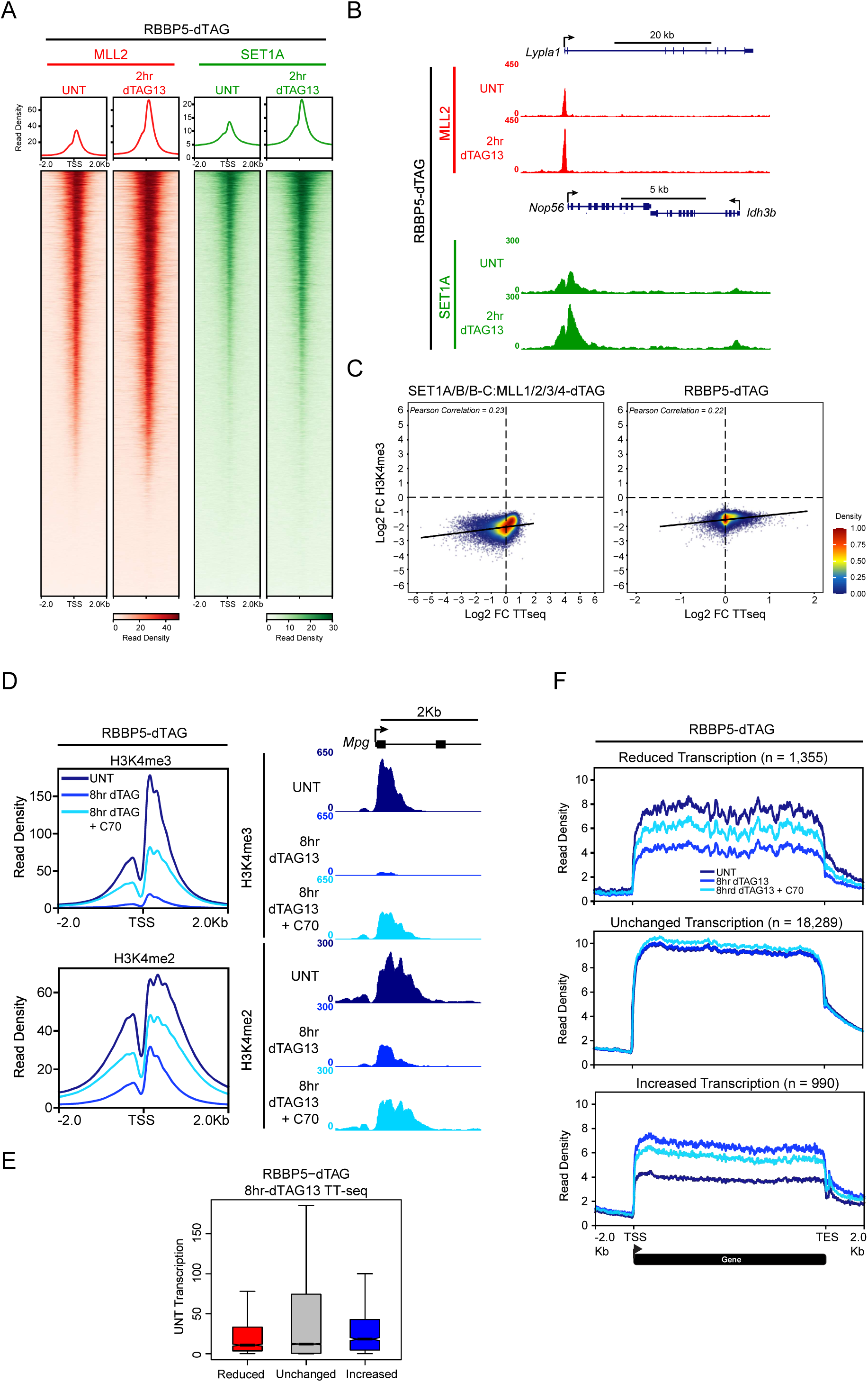
Supplementary SET1/MLL complexes primarily control transcription independently of H3K4me3. A) Metaplots and heatmaps showing MLL2 and SET1A ChIP-seq signal at all TSSs (n = 20,634) after 2 hours of RBBP5-depletion. Heatmaps are sorted in descending order by the untreated ChIP-seq signal. B) Genome coverage tracks showing MLL2 and SET1A ChIP-seq signal at a representative gene (*Lypla1* for MLL2, *Nop56* for SET1A). C) Scatterplots comparing the log2 fold change in H3K4me3 ChIP-seq signal and TT-seq signal in the SET1A/B/B-C:MLL1/2/3/4-dTAG (left) and RBBP5-dTAG (right) cell lines. Only genes with an H3K4me3-associated promoter (n=14,344) are plotted. A 2D density plot overlaying the data is used to illustrate the distribution of the data, with a scale bar showing the colour coding. A linear regression line is plotted, and the Pearson correlation value is also displayed. D) Metaplots of average H3K4me2/3 ChIP-seq signal at all TSSs (n=20,634) after 8 hours of RBBP5-depletion, with or without KDM5 inhibition with C70. Also shown are genome coverage tracks for H3K4me2/3 at a representative gene (*Mpg*) after 8 hours of RBBP5-depletion, with or without KDM5 inhibition with C70. E) Boxplot showing transcription levels (normalised read counts) in untreated RBBP5-dTAG cells at genes showing reduced (n=1,355), unchanged (n=18,289), and increased (n=990) transcription after 8 hours of RBBP5-depletion. F) Metaplots of average TT-seq signal at genes described in (E) after 8 hours of RBBP5-depletion, with or without KDM5 inhibition with C70.

**Figure S6:**
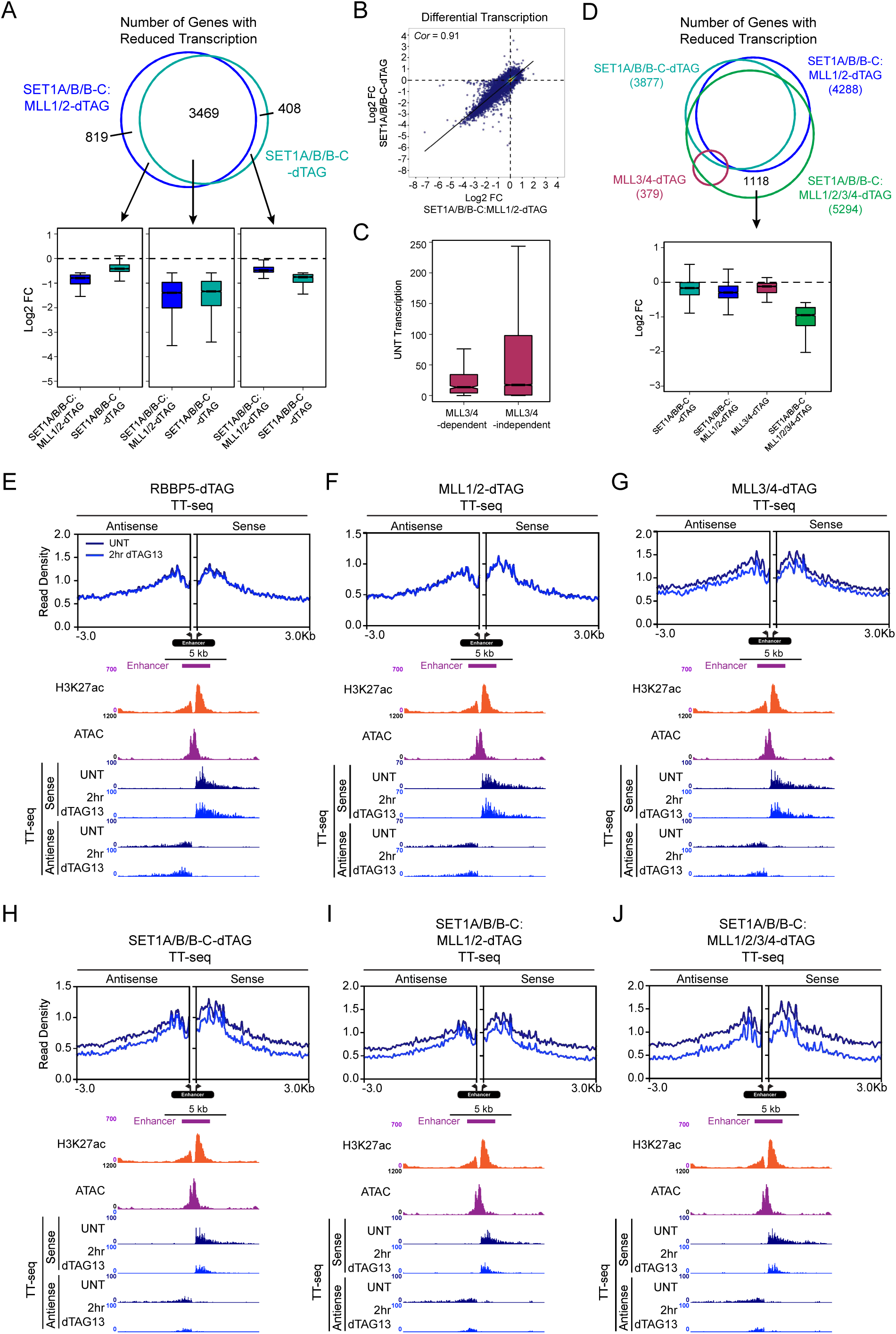
Supplementary to SET1A/B largely explain SET1/MLL complex dependent effects on transcription. A) Venn diagram illustrating overlap between genes reduced in transcription after 2 hours of dTAG13 treatment in the SET1A/B/B-C-dTAG and SET1A/B/B-C:MLL1/2-dTAG cell lines. Shown underneath are boxplots illustrating the log2 fold changes between dTAG13-treated and untreated SET1A/B/B-C-dTAG and SET1A/B/B-C:MLL1/2-dTAG cells at genes reduced in transcription in either or both cell lines. B) Scatterplot comparing TT-seq log2 fold changes between dTAG13-treated and untreated SET1A/B/B-C-dTAG and SET1A/B/B-C:MLL1/2-dTAG cells at all genes (n=20,634). C) Boxplots showing levels of transcription (normalised TT-seq read counts) of MLL3/4-dependent (n=379) and MLL3/4-independent (n=20,254) genes in untreated cells (UNT transcription). D) An Euler plot showing overlap between genes which are reduced in transcription after 2 hour dTAG13 treatment in the indicated cell lines. Shown below is a boxplot comparing TT-seq log2 fold changes between dTAG13-treated and untreated conditions in the indicated cell lines at the genes which are uniquely reduced in transcription in the SET1A/B/B-C:MLL1/2/3/4-dTAG cell line (n=1118). E) Metaplots of average of TT-seq signal at intergenic enhancers (n=3,182) after 2 hours of dTAG13 treatment in the RBBP5-dTAG cell line. Shown below are genome coverage tracks of H3K27ac ChIP-seq and ATAC-seq in untreated E14 mESCs (from Fursova et al. 2021^87^), along with TT-seq in the RBBP5-dTAG cell line after 2 hours of dTAG13 treatment at a representative enhancer (chr15: 96,708,943 – 96,711,064). F) As per (E) but for the MLL1/2-dTAG cell line. G) As per (E) but for the MLL3/4-dTAG cell line. H) As per (E) but for the SET1A/B/B-C-dTAG cell line. I) As per (E) but for the SET1A/B/B-C:MLL1/2-dTAG cell line. J) As per (E) but for the SET1A/B/B-C:MLL1/2/3/4-dTAG cell line.

**Figure S7:**
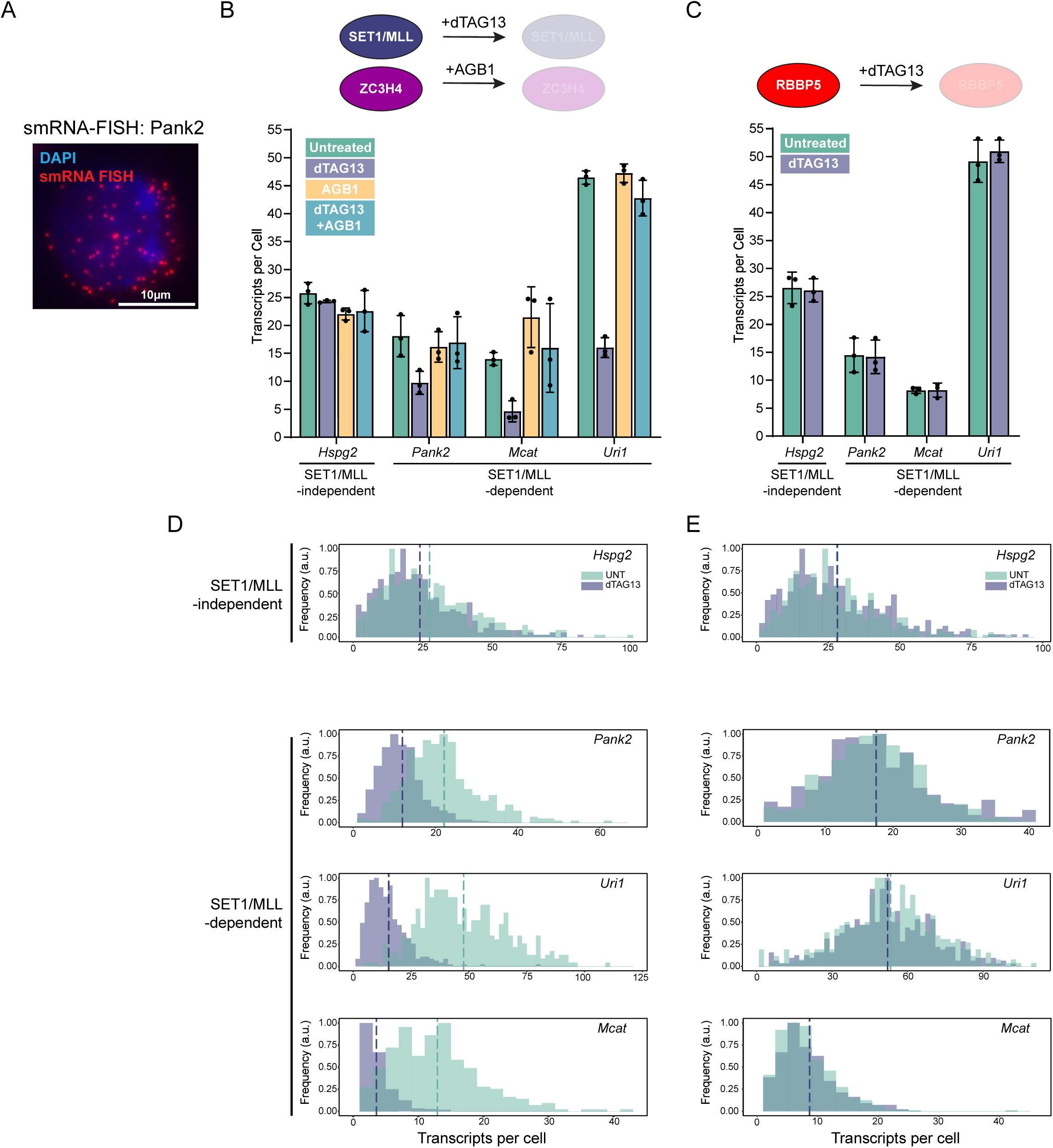
Supplementary to SET1/MLL complexes are essential to sustain transcription ON-period size. A) Representative image of a cell showing single molecule RNA-FISH for *Pank2*. DAPI is shown in blue and individual *Pank2* transcripts are shown in red. B) Bar graphs showing the average number of transcripts per cell from 3 biological replicates indicated as individual dots. Single-cell gene expression profiles for *Hspg2*, *Pank2*, *Mcat*, and *Uri1* after 4 hours of SET1/MLL-depletion (dTAG13), ZC3H4-depletion (AGB1), and SET1/MLL/ZC3H4-depletion (dTAG13+AGB1). At least 283 cells were measured per condition for each replicate. Error bars represent the standard error of the mean. C) Bar graphs showing the average transcripts per cell from 3 biological replicates for *Hspg2*, *Pank2*, *Mcat*, and *Uri1* after 4 hours of RBBP5-depletion (dTAG13). D) Histograms showing the transcript-per-cell distributions for untreated cells (UNT) and cells after 4 hours of SET1/MLL-depletion (dTAG13) for representative individual biological replicates. Vertical dashed lines correspond to average transcript per cell value for the respective condition. The number of measured cells for *Hspg2*, *Pank2*, *Mcat*, and *Uri1* was, respectively: UNT, n=456, n=474, n=312, n=474 dTAG13: n=633, n=400, n=382, n=400. E) Same as D) but after RBBP5 depletion. The number of measured cells for *Hspg2*, *Pank2*, *Mcat*, and *Uri1* was, respectively: UNT, n=465, n=452, n=594, n= 452, dTAG13, n=413, n=590, n=441, n=590.

**Figure S8:**
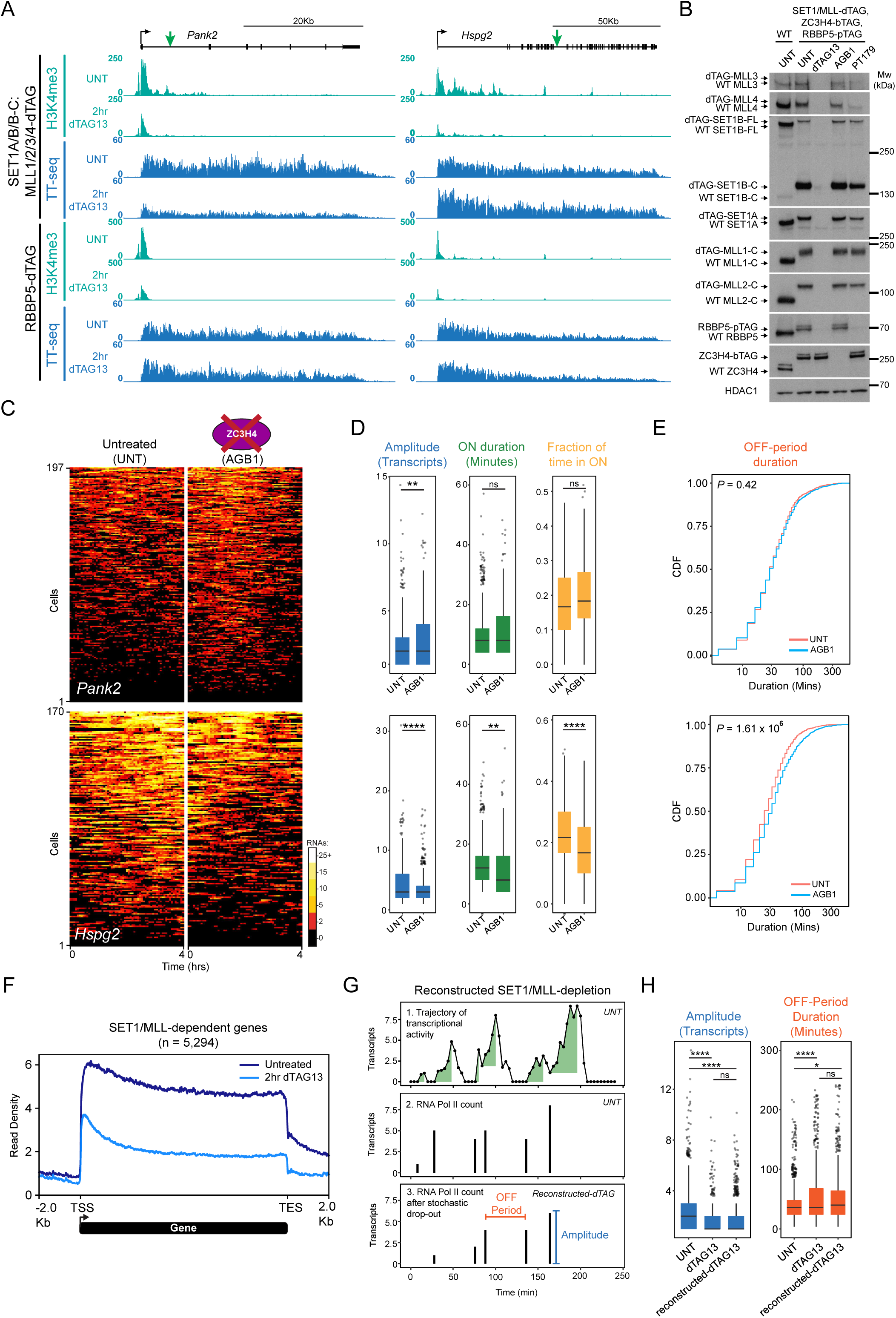
SET1/MLL complexes are essential to sustain transcription ON-period size. A) Genome coverage tracks of TT-seq and H3K4me3 after SET1/MLL-depletion (top) or RBBP5-depletion (bottom) at *Pank2* and *Hspg2*. Green arrows indicate the insertion site of the MS2 arrays in the intron 1 of *Pank2* and the intron 33 of *Hspg2*. B) Western blot showing depletion of SET1/MLL proteins after 2 hours of dTAG13 treatment, depletion of ZC3H4 after 2 hours of AGB1 treatment, and depletion of RBBP5 after 2 hours of PT179 treatment. HDAC1 was used as a loading control. C) Kympographs of transcription trajectories from individual cells for *Pank2* and *Hspg2* after ZC3H4 depletion (AGB1). The number of cells imaged for each gene is indicated on the y-axis, and the duration of the imaging is shown on the x-axis. The amplitude of transcription is colour-coded and shown on the scale bar. 3 biological replicates were imaged for each gene. D) Boxplots showing the distribution of the ON-period amplitude, ON-period duration, and the fraction of time spent in the ON-period for *Pank2* and *Hspg2*. Each data point corresponds to an individual ON-period. Boxes represent the interquartile range (IQR) centred on the median value, with whiskers showing 1.5× IQR and outliers as dots. *P* values were calculated using a two-sided Kolmogorov–Smirnov test, and significant *P* values are shown as asterisks: * p ≤ 0.05, ** p ≤ 0.01, *** p ≤ 0.001, and **** p ≤ 0.0001. n.s. corresponds to p > 0.05. E) Cumulative distribution function plots of the OFF-period duration after ZC3H4-depletion. F) Metaplot of average TT-seq signal at all SET1/MLL-dependent genes (n = 5,294) after SET1/MLL-depletion. G) Cartoon illustrating how dTAG13 live-cell transcription trajectories were reconstructed from UNT data. (top) Representative transcription trajectory illustrating the reconstruction of the effect of SET1/MLL-depletion on ON-period amplitude and OFF-period duration from experimental data from untreated cells. Each trajectory of transcriptional activity is simplified into RNA Pol II counts (middle), which then were randomly subsampled to reconstruct the effect of a stochastic drop out of RNA Pol IIs following SET1/MLL-depletion (bottom). Note, this primarily affects ON-period amplitude but may also affect OFF-period duration at instances whenever ON-periods are entirely terminated. H) Boxplots showing the distribution of the ON-period amplitude and OFF-period duration. The reconstructed SET1/MLL-depletion data is plotted next to the experimental data for the untreated and SET1/MLL-depletion conditions. Each data point corresponds to an individual ON-period. Boxes represent the interquartile range (IQR) centred on the median value, with whiskers showing 1.5× IQR and outliers as dots. *P* values correspond to two-sided Kolmogorov–Smirnov test, and significant *P* values are shown as asterisks: * p ≤ 0.05, ** p ≤ 0.01, *** p ≤ 0.001, and **** p ≤ 0.0001. n.s. corresponds to p > 0.05.

